# A *Salmonella* type I toxin promotes systemic infection by inhibiting F_o_F_1_ ATP synthase

**DOI:** 10.1101/2025.05.26.656083

**Authors:** Seoyeon Kim, Seonggyu Kim, Eun-Jin Lee

## Abstract

Intracellular pathogens including *Salmonella* Typhimurium survive within macrophage phagosomes by forming non-replicating persisters. Toxins from toxin-antitoxin systems have been implicated in persister formation as key contributors, although the biological targets of toxins remain largely unknown. Here, we report that *Salmonella* IbsA, a 19-amino-acid-long type I toxin, induces growth arrest and decreases intracellular ATP levels. An unbiased large-scale bacterial two-hybrid screen identified 54 IbsA-interacting targets, including proteins involved in oxidative phosphorylation and cell division. Among these, IbsA toxin specifically targets two subunits of the bacterial F_o_F_1_ ATP synthase. IbsA interacts with the *a* and *b* subunits in the membrane-bound F_o_ sector and inhibits proton translocation, thereby decreasing ATP levels. In turn, this IbsA-mediated decrease facilitates *Salmonella* spread between macrophages, promoting systemic infection. Accordingly, *ibsA* deletion decreases bacterial burden in organs and attenuates mouse virulence. Therefore, unlike other bacterial toxins, the IbsA toxin enhances *Salmonella* pathogenicity by decreasing the bacterium’s ATP levels.

## INTRODUCTION

Intracellular pathogens, including *Salmonella* enterica serovar Typhimurium, are capable of surviving and replicating within macrophage vacuole (1, 2). Despite the fact that intracellular bacteria within the vacuole are protected from immune responses, they have to deal with stressful conditions (3). For successful survival, *Salmonella* enterica has evolved several mechanisms to survive within rapidly changing and stressful conditions during infection (4). One such survival mechanism is the formation of bacterial persisters, which is a non-replicating, multidrug-tolerant subpopulation that arises from phenotype switching (5, 6). The toxin-antitoxin (TA) system has been suggested as a genetic component involved in bacterial persister formation inside host cells as well as in response to antibiotic exposure (7–9). The TA system is a two-component genetic module, which consists of a toxin targeting vital cellular processes and its cognate antitoxin antagonizing toxin’s activity. TA system loci were initially recognized on plasmid R1 that encodes *hok*/*sok,* a module involved in post-segregational killing (PSK), which kills cells upon plasmid loss (10, 11). Later, many TA modules were discovered on bacterial chromosomes. Further bioinformatics analyses revealed that TA systems are widely distributed in bacteria and archaea, mainly in prokaryotes (12, 13). Toxins in TA systems are typically diverse proteins that hinder key cellular activities such as cell proliferation and replication, whereas antitoxins are either RNA or protein that neutralize the activity of their cognate toxin. Currently, TA systems are classified into eight types based on the nature of the antitoxin or the mechanism by which the antitoxin neutralizes the toxin (8, 14). Among these, type I TA systems contain a noncoding RNA antitoxin that base-pairs with toxin mRNA and prevents protein synthesis (8). Antitoxin RNAs in type I TA modules can be further divided into two groups: *cis*-encoded small RNAs (sRNAs) overlapping with their complementary toxin mRNA, and *trans*-encoded sRNAs located at divergent chromosomal positions (15). These antitoxin sRNAs inhibit toxin expression by blocking the Shine-Dalgarno sequence of toxin mRNAs, inducing toxin mRNA cleavage, or binding to upstream regions to compete with ribosome loading (16–19). Although the regulatory mechanisms of antitoxin sRNAs are relatively well studied, little is known about the function and biological targets of type I toxins. With the exceptions of SymE encoding an endoribonuclease and RalR encoding a DNase, most type I toxins are small hydrophobic proteins that contain a single α-helical transmembrane domain (15, 16). Overexpression of small membrane toxins including Hok, TisB, Ldr, and Ibs appear to be involved in membrane depolarization and ATP loss (20–29). In the case of Hok, it has been reported that Hok oligomerizes in the membrane and induces the pore formation, thereby causing ATP leakage (30). ATP depletion via pore formation is widely accepted as a general mechanism of small hydrophobic toxins due to their similarities in length, hydrophobic nature, and ATP-depleting effect (20–24). In the intracellular pathogen *Salmonella* Typhimurium SL1344, five type I TA modules-*hok/sok*, *symER*, *tisB/istR, ldr/rdl*, and *ibs/sib*-have been identified (31). However, the molecular mechanisms by which these toxins achieve non-replicating state remains largely unknown. In this study, we sought to identify the biological targets of a *Salmonella* type I toxin to understand its physiological function(s).

Among type I TA system toxins, Ibs family toxins were initially discovered while examining Sib antitoxin, a *cis*-encoded small RNA (sRNA) in *Escherichia coli* K-12 (29). In *E. coli*, five *ibs*/*sib* pairs exist, and in most cases-except for IbsA-the repression of cognate toxin by antitoxin is highly selective due to the *cis*-encoded nature of target recognition domains in Sib RNA (29, 32). Similar to other type I toxins, Ibs toxins are small hydrophobic membrane proteins and their overexpression rapidly depolarizes the membrane and prevents colony formation on solid medium (29). Additionally, it has been reported that highly hydrophobic residues in the core of Ibs toxin proteins are required for toxicity (33). However, it is unclear how Ibs proteins function as toxins, as little is known about the biological targets of type I toxins. Here, we report the characterization of the STM_3875 gene as the IbsA toxin from *Salmonella* enterica serovar Typhimurium 14028s and the identification of IbsA target proteins using genome-wide bacterial two-hybrid screening. We demonstrate that IbsA toxin interacts with the F_o_F_1_ ATP synthase and inhibits proton translocation-coupled ATP synthesis. Contrary to the classical idea that toxins are involved in persister formation by promoting a non-replicating state and contributing to non-responsiveness to host immune systems or antibiotic treatment, physiologically expressed IbsA toxin mediates ATP depletion via the F_o_F_1_ ATP synthase, thereby enabling *Salmonella* to promote transmission between macrophages and establish systemic infection. Therefore, these findings reinforce the notion that the function of bacterial toxins must be examined within their biological context.

## RESULTS

### The *ibsA* gene encodes a 19 amino acid-long type I toxin in *Salmonella*

To search for a type I toxin in *Salmonella* Typhimurium 14028s, we referred to a previous report on *Escherichia coli* (29) and analyzed an intergenic sequence between the *yqiK* gene and *rfaE* gene encoding an inner membrane protein and lipopolysaccharide biosynthesis protein RfaE, respectively. Unlike *E. coli* MG1655 that contains two type I toxin genes, *ibsD* and *ibsE* (29), the intergenic region of *Salmonella* Typhimurium contains a single open reading frame, STM14_3875 encoding a 65 aa long protein with a C-terminal transmembrane domain (Fig. 1a). Following the annotation in *Salmonella* Typhimurium SL1344, we designated STM14_3875 as *ibsA* (31). The *Salmonella ibsA* gene has two possible start codons; upstream GTG codon and downstream ATG codon. The upstream GTG codon produces a 65 aa long protein, whereas the downstream ATG codon produces a 19 aa long protein, which is similar to other Ibs toxins in *E. coli* (Fig. 1b, Fig. 1c). To examine which start codon produces type I toxin protein, we cloned DNA sequences corresponding to either a longer GTG-started *ibsA* gene or a shorter ATG-started *ibsA* gene into a plasmid, which then will be fused to a C-terminal 8×Myc tag. Interestingly, although GTG-started *ibsA* mRNA was longer than ATG-started *ibsA* mRNA (Fig. 1d), no difference in protein size was detected between the two constructs (Fig. 1e), suggesting that the IbsA protein is translated from the internal ATG codon, producing a 19 aa long protein.

**Fig. 1.**
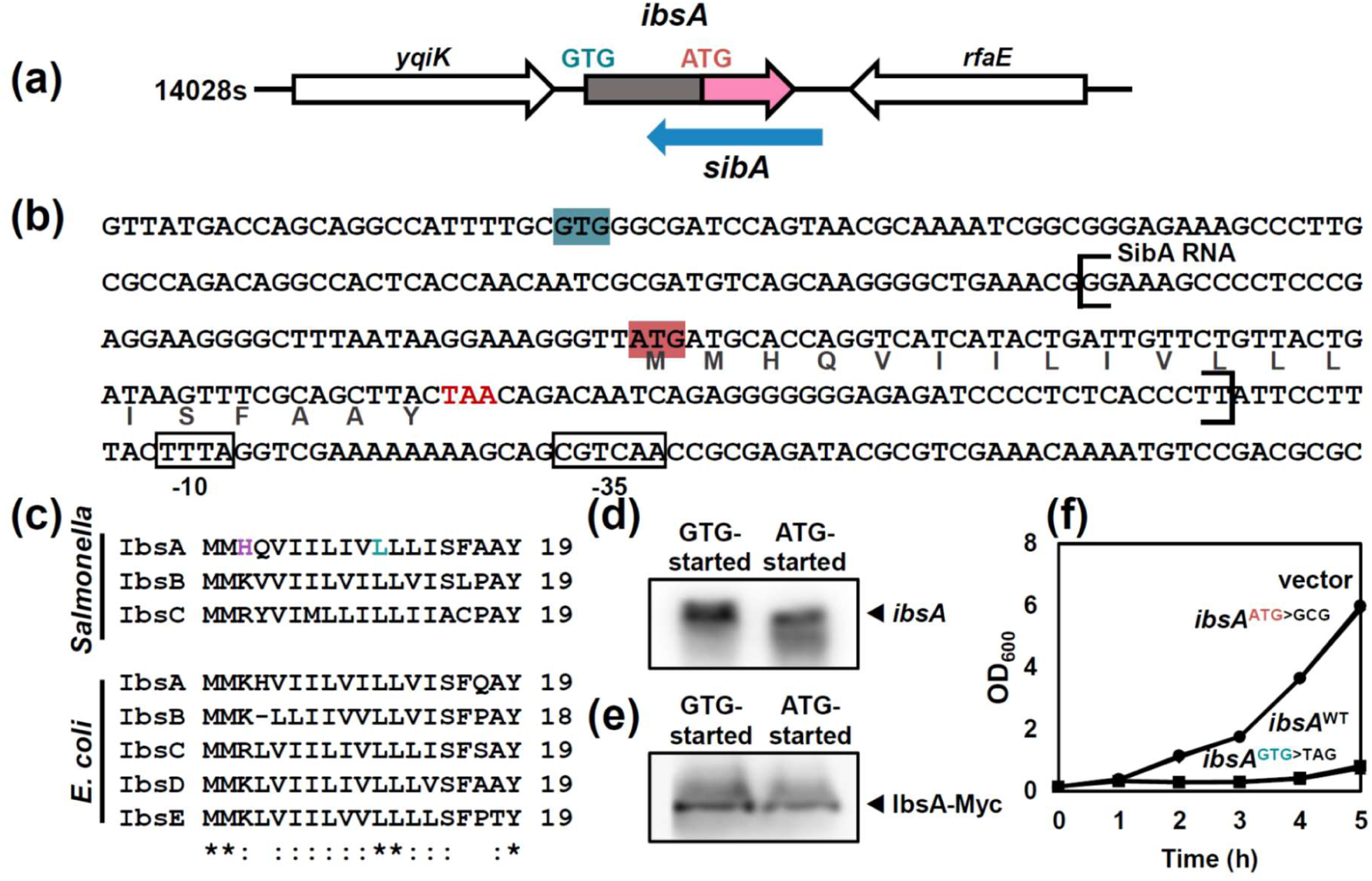
*ibsA* gene encodes type I toxin protein in *Salmonella* Typhimurium (a) Genetic map of the *ibsA* gene locus in *Salmonella* Typhimurium 14028s. (b) DNA sequences of *ibsA* chromosomal region. Two possible start codons in the gene are highlighted in red and blue. *sibA* sequence is indicated by brackets and promoter elements of *sibA* are indicated by boxes. (c) Amino acid sequences of Ibs toxins from *Salmonella enterica* serovar Typhimurium 14028s and *E. coli* MG1655. His3 and Leu11 residues in *Salmonella* IbsA are indicated in purple and cyan. (d, e) Northern blot analysis of *ibsA* mRNAs (d) and Western blot analysis of IbsA-Myc proteins (e). Total RNA and cell lysates were extracted from *Salmonella* harboring pBAD33 plasmid with either GTG-started or ATG-started *ibsA-*8xMyc. Cells were grown in N-minimal medium containing 10 mM Mg^2+^ for 2 h after addition of 10 mM arabinose. (d) RNA samples (10 µg) were analyzed by Northern hybridization using oligonucleotide probe specific to Myc. (e) Cell lysates were analyzed by immunoblot assay using anti-Myc antibodies. (f) Growth curves of *Salmonella* harboring pBAD33 (●), pBAD33-*ibsA* (▪), pBAD33-*ibsA*^GTG>TAG^ (▴) or pBAD33-*ibsA*^ATG>GCG^ (◆). The strains were grown at 37°C in LB medium and *ibsA* was induced by addition of 10 mM arabinose when the cell reached at OD_600_=0.1.

To further clarify, we cloned the full *ibsA* region containing both GTG and ATG start codons into an arabinose-inducible plasmid and measured growth upon arabinose induction (Fig. 1f). Induction of the long *ibsA* region arrested *Salmonella* growth compared to the empty vector, indicating that this region indeed produces a toxin (Fig. 1f). Substitution of the internal ATG start codon with a GCG alanine codon completely abolished toxin activity, whereas substitution of the upstream GTG start codon with a TAG stop codon still retained toxin activity (Fig. 1f). Therefore, *Salmonella ibsA* gene is misannotated and, like other *ibs* toxin genes, it is 60 nt in length, encoding a 19 amino acid-long protein.

### His3 and Leu11 residues are required for the bacteriostatic activity of IbsA toxin

IbsA is a hydrophobic, single-pass transmembrane protein with its N-terminus exposed to the cytoplasm (Fig. 2a). To identify key residues for IbsA toxin activity, alanine scanning mutagenesis was conducted in the 19 aa version of the IbsA toxin. Similar to Fig. 1f, *Salmonella* expressing 19 aa wild-type IbsA attenuated growth compared to the empty vector (Fig. 2b). Unlike wild-type IbsA, the H3A and L11A substitutions restored growth to levels comparable to the empty vector in LB medium (Fig. 2b), indicating that the His3 and Leu11 residues are functionally required for IbsA toxin activity. As controls, other substitutions still retained growth inhibitory activity of IbsA comparable to the wild-type IbsA (Fig. 2b). Ibs proteins are conserved in enteric bacteria including *E. coli*, *Salmonella* Typhimurium, *Shigella flexneri,* and *Citrobacter koseri* (29). It is interesting to note that, when we compared IbsA-like toxins in *S*. Typhimurium 14028s and *E. coli* MG1655, all of the 3^rd^ residues in IbsA-like toxins are positively charged amino acids such as His, Lys, or Arg and the 11^th^ residues are leucine (Fig. 1c). In addition, when IbsA toxin protein structure was predicted using AlphaFold3 server (34), both His3 and Leu11 residues appeared to be located at the same side within the transmembrane helix (Fig. 2a).

**Fig. 2.**
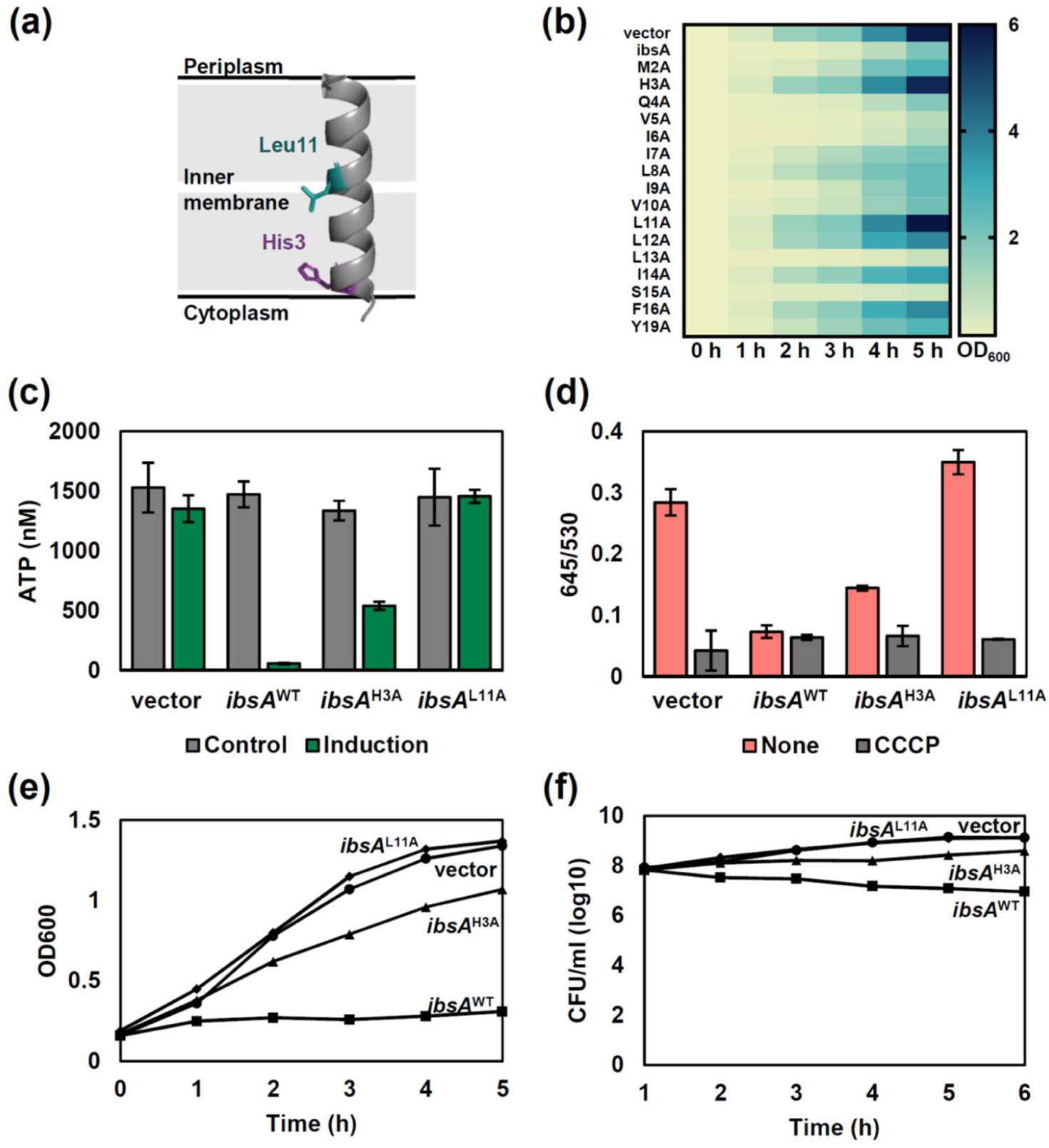
His3 and Leu11 of the IbsA toxin are essential for bacteriostatic activity (a) A modeled structure of the IbsA protein predicted by AlphaFold. The His3 residue is indicated with purple stick and Leu11 residue is indicated with cyan stick. (b) Histogram of growth curves of alanine scanning variants of IbsA. *Salmonella* harboring pBAD33 carrying *ibsA* wild-type or its derivatives were grown at 37 °C in LB medium and the expression of *ibsA* was induced by addition of 10 mM arabinose when the cell reached at OD_600_=0.1. (c) Intracellular ATP levels of *Salmonella* wild-type strains harboring an arabinose-inducible plasmid with the wild-type *ibsA*, *ibsA*^H3A^, or *ibsA*^L11A^ genes, and the empty vector in the presence (green) or absence (grey) of arabinose. The strains were grown at 37°C in N-minimal medium containing 10 mM Mg^2+^ and the expression of *ibsA* was induced by addition of 10 mM arabinose when the cell reached at OD_600_=0.1. The cells were grown for additional 2 h and harvested for measurement of ATP level. (d) Membrane potential of *Salmonella* wild-type strains harboring either empty vector or pBAD33-*ibsA*, *ibsA*^H3A^, *ibsA*^L11A^. Bacteria were grown as described in (c). Cells were harvested and incubated with membrane potential indicator dye DiOC_2_ in either the presence or absence of the proton ionophore CCCP. After 30 min, red/green fluorescence ratio was determined. (e) H3A and L11A substitutions restores growth similar to that of the empty vector. Wild-type harboring either empty vector (●) or pBAD33-*ibsA* (▪), *ibsA*^H3A^ (▴), *ibsA*^L11A^ (◆) were grown at 37°C in N-minimal medium containing 10 mM Mg^2+^ and the expression of *ibsA* was induced by addition of 10 mM arabinose when the cell reached at OD_600_=0.1. The absorbance at OD_600_ was measured every hour. (f) IbsA toxin’s effect is bacteriostatic, not bactericidal. Bacteria were grown as described in (e). Cells were diluted and plated on LB agar plates every hour.

Type I toxins were proposed to target the bacterial membrane, which lead to membrane depolarization and ATP depletion (24). To address whether IbsA acts in a similar manner, we measured intracellular ATP levels upon *ibsA* induction (Fig. 2c). Intracellular ATP levels of *Salmonella* expressing IbsA dramatically decreased compared to those expressing empty vector (Fig. 2c), which can affect bacterial growth. Likewise, *ibsA* induction also decreased fluorescence from membrane potential-dependent dye 3,3′-diethyloxacarbocyanine iodide (DiOC_2_), similar to when we treated with protonophore CCCP (carbonyl cyanide 3-chlorophenylhydrazone)(Fig. 2d), suggesting that IbsA toxin arrests bacterial growth by decreasing intracellular ATP levels and membrane potential. As controls, non-induction control or the *ibsA* variants with the L11A substitution remained high levels of ATP and membrane potential and the *ibsA* H3A substitution did not efficiently decrease ATP levels and membrane potential (Fig. 2c and 2d).

We wondered whether the IbsA-dependent growth inhibiton is bacteriostatic or bactericidal. To test this, we measured optical density and colony-forming unit (CFU) of *Salmonella* strains expressing wild-type IbsA or its derivatives (Fig. 2e and 2f). Although wild-type IbsA overexpression arrested bacterial growth, CFUs of the strain expressing wild-type IbsA were comparable to those expressing vector or IbsA derivatives with the H3A or L11A substitutions (Fig. 2f). These data indicate that IbsA toxin activity is bacteriostatic, not bactericidal.

Toxin-antitoxin systems are involved in persister formation, which a fraction of cells enters a dormant state and exhibits antibiotic tolerance (15). Among type I toxins, TisB was previously reported to increase the formation of persister cells tolerant to ciprofloxacin, an antibiotic inhibiting gyrase and inducing DNA damage (35, 36). We measured whether IbsA can induce antibiotic-induced persister formation (Fig. S1a).

When we treated ciprofloxacin, IbsA-expressing *Salmonella* increased persister formation by ∼10^2^ fold compared to those expressing the empty vector or IbsA variants with H3A or L11A substitutions (Fig. S1b). This indicates that IbsA overexpression promotes antibiotic-induced persister formation after ciprofloxacin treatment.

### IbsA toxin’s effect is carbon source-dependent

Interestingly, we noticed that the decrease in intracellular ATP levels in IbsA-overexpressing *Salmonella* was not detected when we switched carbon source in the N-minimal medium from glycerol to glucose (Fig. 3a-3c). Given that glucose enters at the top of the glycolytic pathway and is expected to yield higher ATP levels than glycerol, this result suggests that high levels of ATP production overcome the effect of IbsA toxin. To explore this further, we then selected another carbon source, succinate, which enters in the middle of the tricarboxylic acid cycle, to see whether IbsA’s effect on ATP levels would be maintained (Fig. 3a). In succinate-containing medium, IbsA expression decreased ATP levels similarly to glycerol-containing medium (Fig. 3d). These data indicate that the effect of IbsA on ATP levels is dependent on the carbon source. As controls, strains expressing the vector or IbsA variants did not exhibit such decreases in ATP levels under all tested conditions (Fig. 3b-3d).

**Fig. 3.**
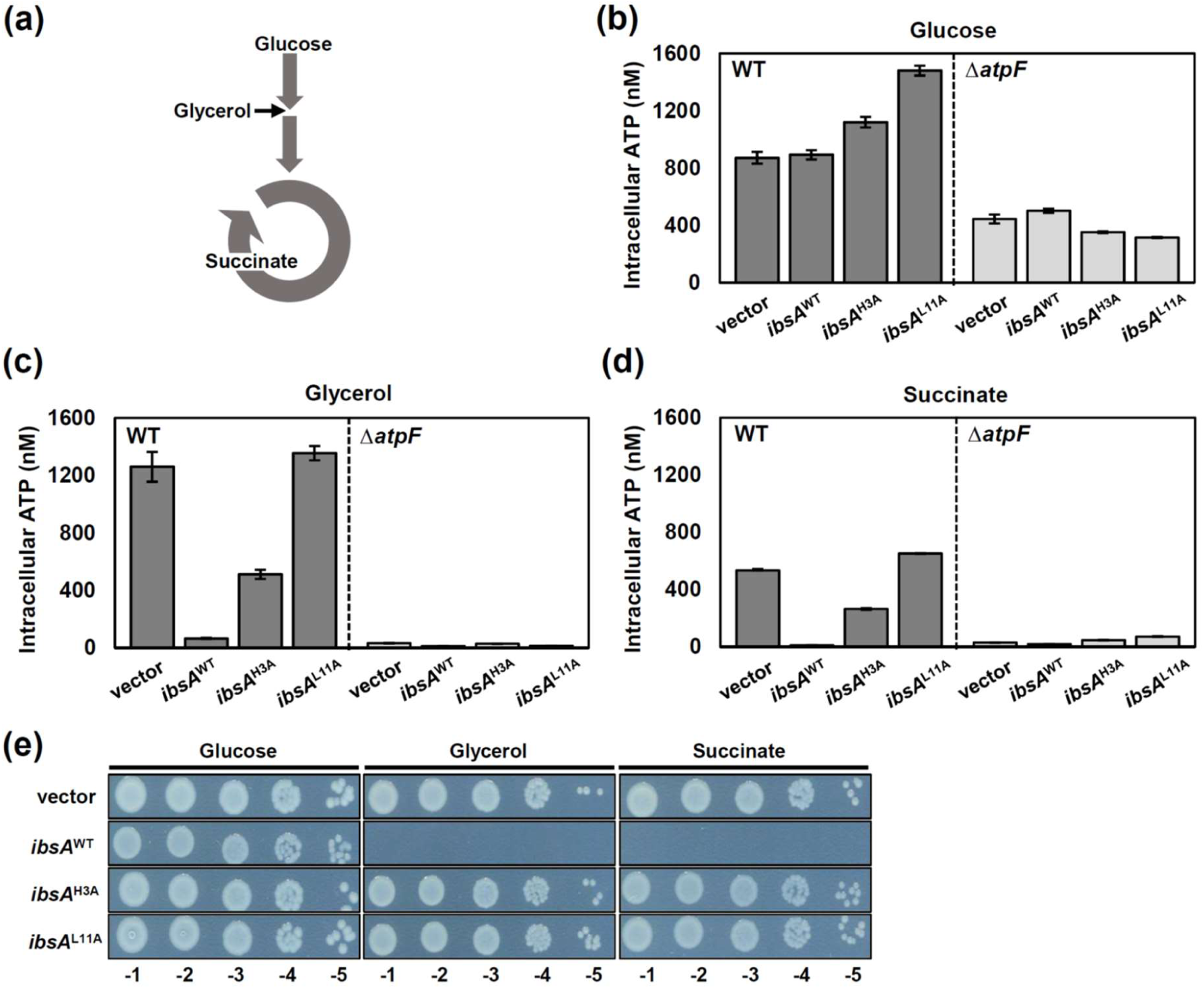
IbsA toxin decreases intracellular ATP levels and colony formation in glycerol- or succinate-containing media but not in glucose-containing media (a) Schematic cartoon of metabolic pathways in bacteria with carbon sources used in this study. (b-d) Intracellular ATP levels of wild-type or *ΔatpF Salmonella* strains harboring the arabinose-inducible plasmid with wild-type *ibsA*, *ibsA*^H3A^, or *ibsA*^L11A^ genes, and the empty vector. Cells were grown at 37°C in N-minimal medium containing 0.2% glucose (b), 0.2% glycerol (c), or 0.2% succinate (d) as a sole carbon source and *ibsA* expression was induced by adding 10 mM arabinose when the cell reached at OD_600_ of 0.1. The cells were grown for additional 2 hours and harvested for measuring ATP levels. (e) Dilution spotting on N-minimal solid media with different carbon sources. 14028s harboring either empty vector or pBAD33-*ibsA*, *ibsA*^H3A^, or *ibsA*^L11A^ were serial diluted and spotted on N-minimal agar plate containing 10 mM arabinose with either 0.2% glucose, glycerol or succinate.

The carbon source-dependent effect of IbsA toxin was more exaggerated when we performed spot dilution assays on solid media with different carbon sources. Similarly to what we observed in ATP levels, IbsA-expressing cells did not form colonies on solid minimal media containing glycerol or succinate (Fig. 3e). However, IbsA expression did not exhibit any defect in colony formation in glucose-containing media, similar to the vector or the IbsA inactive variants (Fig. 3e). These results indicate that IbsA toxin’s effect differs depending on the ATP levels produced under given conditions.

### IbsA toxin interacts with the *a* and *b* subunits in the membrane-embedded F_o_ complex of ATP synthase

Previously, HokB was proposed to function as a type I toxin via a mechanism by which it targets the bacterial membrane, oligomerizes, and induces pore formation, leading to membrane disruption and ATP loss (30). In this case, a single cysteine residue located in the periplasm was identified to be essential for oligomerization and dynamic pore formation (37). Although IbsA also decreased ATP levels and membrane potential (Fig. 2), we suspected that IbsA might function differently because: i) IbsA toxin is shorter in length and thus lacks a periplasmic region, ii) IbsA does not contain a cysteine residue required for disulfide bond formation, iii) the carbon-source dependent effect of IbsA toxin reflects that IbsA may target metabolic pathways other than pore formation, and iv) when we measured extracellular ATP levels in supernatants, extracellular ATP levels were relatively low and did not correlate with IbsA toxin’s activity (Fig. S2).

Therefore, we set up a screening experiment to identify IbsA-interacting proteins. To explore this, a bacterial adenylate cyclase two-hybrid (BACTH) assay was employed to screen *Salmonella* gDNA library (Fig. 4a). The *ibsA* gene was C-terminally fused to the T25 domain of adenylate cyclase, and enzyme-digested or mechanically fragmented gDNA libraries were N- or C-terminally fused to the T18 domain. In principle, if T25-fused IbsA physically interacts with a T18-fused target protein translated from gDNA fragments, these two domains reconstitute a functional adenylate cyclase to synthesize cAMP and lead to produce *β*-galactosidase, which gives red colonies on MacConkey plate (38). After screening to cover more than 120-fold genome equivalents, we selected 100 positive clones as potential IbsA-interacting targets (Fig. 4b). DNA sequence analysis revealed that these clones include T18-fused DNA fragments corresponding to many genes involved in energy production and oxidative phosphorylation, such as lipid/carbohydrate metabolism, electron transport chain, and ATP synthase, as well as other functional categories (Fig. S3). After reintroducing the original clones, we narrowed down the hits to 54 IbsA-interacting targets and analyzed by their functional categories (Table 1). Most of the IbsA-interacting targets were located in the membrane (Fig. S3b) and were enriched in functions such as energy metabolism and membrane transport (Table 1).

**Fig. 4.**
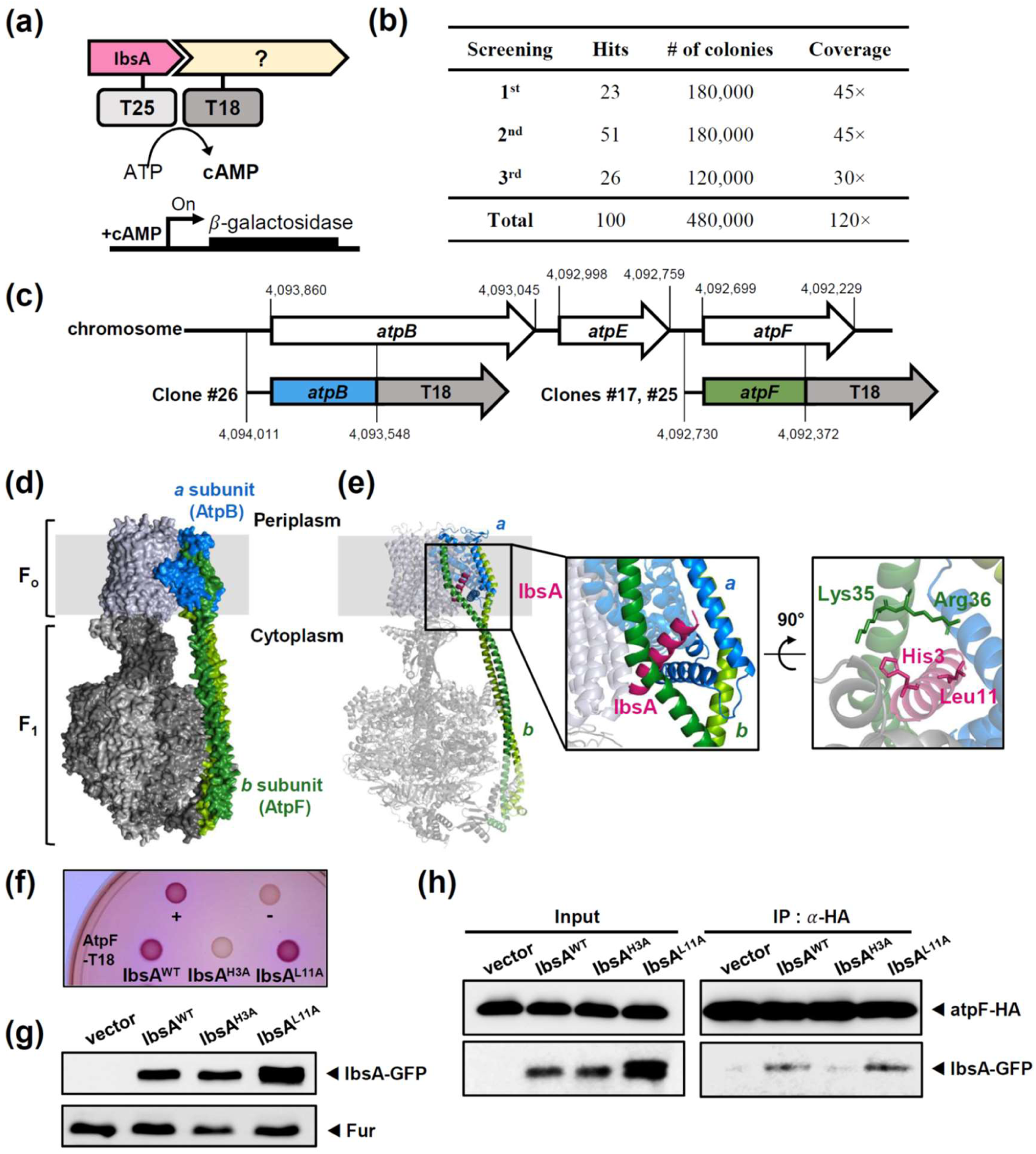
Bacterial two-hybrid screening identifies *a* and *b* subunits of ATP synthase Fo complex as IbsA targets (a) Schematic representation of bacterial two-hybrid (BACTH) screening. The BACTH system is an effective tool for identifying target proteins that interact with the membrane-bound IbsA toxin. The 25 kDa domain (T25) and the 18 kDa domain (T18) of adenylate cyclase are individually fused to either a bait protein (IbsA) or library proteins translated from genomic DNA fragments. Both clones are then co-transformed into a bacteria strain lacking adenylate cyclase. If IbsA and a candidate protein interact, the fused T25 and T18 domains functionally complement adenylate cyclase activity and produce cAMP, thereby activating *β*-galactosidase. Positive clones can be selected using MacConkey media containing maltose. (b) Summary of the genome coverage of the BACTH screening. (c) Genetic map of three DNA fragments isolated from IbsA-interacting clones. In this screen, random genomic DNA fragments were generated by sonication, blunt-ended, and inserted into pUT18, which was digested with SmaI. (d) AlphaFold 3 prediction of *Salmonella* F_o_F_1_ ATP synthase (ipTM=0.68, pTM=0.7). (e) AlphaFold3-assisted prediction of interaction between *Salmonella* F_o_F_1_ ATP synthase and IbsA toxin (ipTM=0.69, pTM=0.71). ATP synthase F_o_ complex subunit *a* (AtpB) is shown in blue and subunit *b* (AtpF) is shown in green. In the *a* and *b* subunits, regions corresponding to the DNA fragments recovered from bacterial two hybrid screening are shown solid blue and green. The possible location of IbsA and key residues are also indicated in the inset. (f) Bacterial two-hybrid assay between IbsA and AtpF. *Escherichia coli* BTH101 strains harboring two plasmids, pUT18 and pKT25 derivatives expressing the N-terminal fusions of the *cyaA* T25 fragments to the coding regions of *ibsA, ibsA*^H3A^, or *ibsA*^L11A^ genes, and a C-terminal fusion of the *cyaA* T18 fragment to the coding region of *atpF*. Cells were spotted onto MacConkey agar plates containing 0.1 mM IPTG and incubated at 30°C for 48 h. Red-colored colonies indicate a positive interaction. (g) Western blot analysis of IbsA and its derivatives cloned in pBAD33 with an N-terminal GFP tag. Cell lysates were analyzed by immunoblot using anti-GFP and anti-Fur antibodies. (h) Anti-HA pull-down assay was performed in an *atpF*-HA tagged strain with pBAD33 harboring *ibsA* and its derivatives fused with an N-terminal GFP tag. For input, crude extracts were detected with anti-HA and anti-GFP antibodies. For immunoprecipitation, the lysed extracts were incubated with anti-HA antibody-coated beads overnight, and eluted fractions were detected with anti-HA and anti-GFP antibodies.

**Table. 1.**
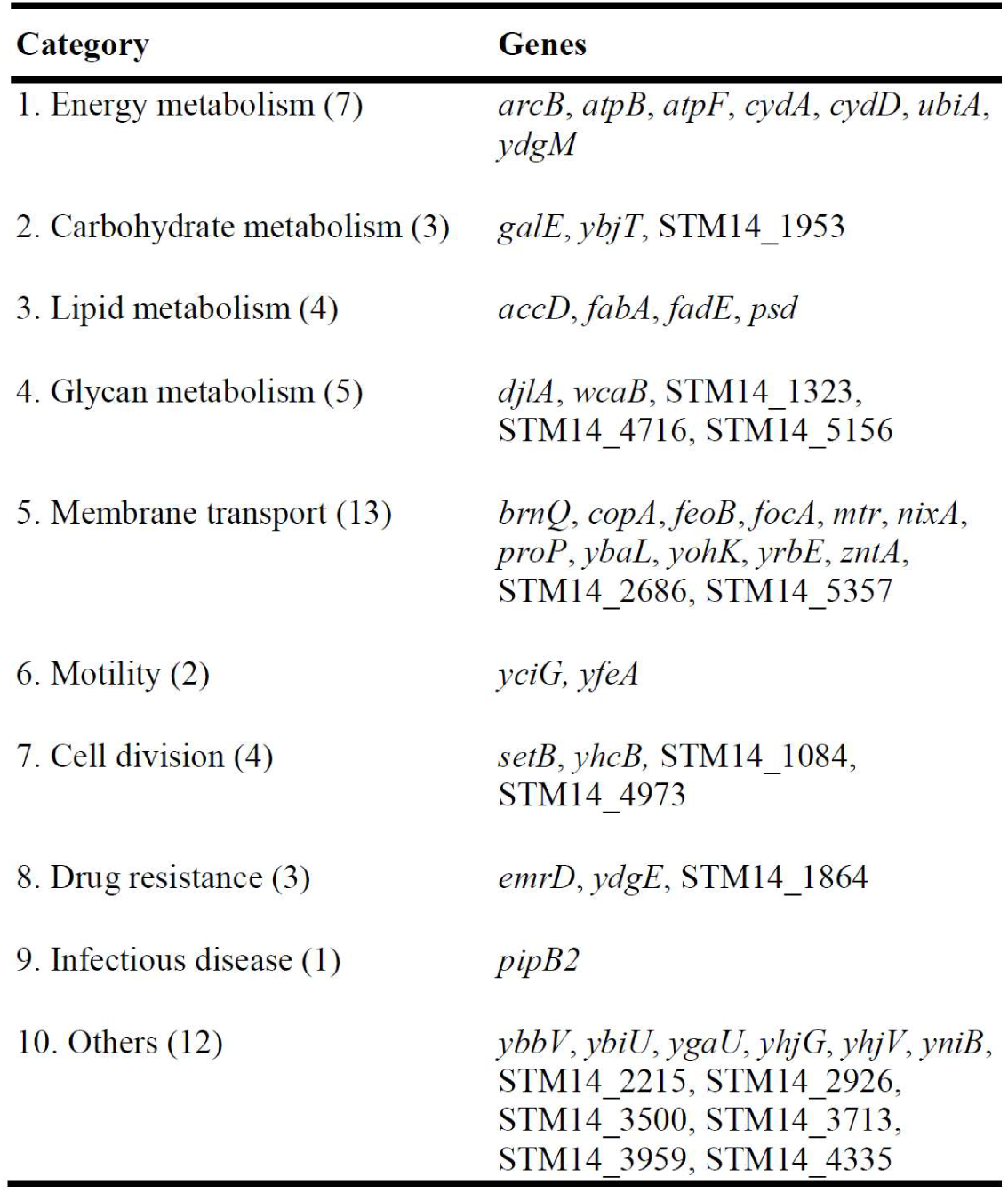
Functional classification of gene fragments of screened IbsA-interacting targets based on annotated functions.

Among the positive hits, we immediately focused on three clones (clones 17, 25, and 26) corresponding to *atpB* and *atpF* genes, which encode the *a* and *b* subunits of F_o_F_1_ ATP synthase, respectively (Fig. 4c), given that IbsA expression decreases intracellular ATP levels and F_o_F_1_ ATP synthase is a multiprotein complex producing most of the ATP in bacteria (39, 40). ATP synthase comprises a membrane-embedded F_o_ complex conducting proton translocation coupled with *c*-ring rotation and a catalytic F_1_ head protruding toward the cytoplasm (41). As both the F_o_ *a* subunit (*atpB*) and *b* subunit (*atpF*) are located in the bacterial inner membrane (42), it makes sense as IbsA-interacting proteins, considering that IbsA toxin is also predicted as a single-helix membrane peptide. An AlphaFold-assisted structural prediction showed that IbsA can bind to the F_o_ *a* subunit (*atpB*) and *b* subunit (*atpF*) near the cytoplasmic surface of the inner membrane (Fig. 4d and 4e). To confirm the interaction between IbsA and the ATP synthase *b* subunit, the full-length *atpF* gene was cloned again into pUT18 and introduced into BTH101 strains expressing T25-IbsA^WT^, or IbsA variants with H3A or L11A substitutions (Fig. 4f). Wild-type IbsA exhibited a red color, indicating that the full-length *b* subunit also interacts with IbsA (Fig. 4f). However, the H3A-substituted IbsA did not produce a red color on MacConkey plates (Fig. 4f), even though the protein levels of IbsA variants were similar to wild-type (Fig. 4g). Similarly, a C-terminally HA-tagged *b* subunit (AtpF-HA) immunoprecipitated an N-terminally GFP-tagged wild-type IbsA toxin but not the H3A-substituted IbsA (Fig. 4h). This is in agreement with the structural prediction that His3 of IbsA faces the *b* subunit (Fig. 4e). In the same prediction, IbsA L11 appears to be close to the *a* subunit but not the *b* subunit.

Consistent with this, the L11A-substituted IbsA still interacted with *b* subunit in both the bacterial two-hybrid and immunoprecipitation assays, although L11 residue is also required for IbsA toxin activity (Fig. 4f and 4h). This supports the conclusion that IbsA interacts with the *b* subunit of F_o_F_1_ ATP synthase, and the His3 residue in IbsA is required for this interaction.

### IbsA toxin inhibits the proton-translocating activity of F_o_F_1_ ATP synthase

In the F_o_F_1_ ATP synthase, the *a* subunit constitutes the interface of proton-translocating channel in the membrane-embedded F_o_ complex, and the *b* subunit links the membrane-bound F_o_ and catalytic F_1_ complexes to couple proton motive force with ATP synthesis (41). Given that IbsA interacts with *a* and *b* subunits in the F_o_ complex of ATP synthase and IbsA expression leads to a decrease in intracellular ATP levels (Fig. 2 and 4), we reasoned that the decrease in ATP levels upon IbsA expression could be due to IbsA interfering with the proton-translocating activity of the F_o_ complex, which is coupled to ATP synthesis in the F_1_ complex. Because the F_o_F_1_ ATP synthase is a reversible enzyme, it can also catalyze a reversible reaction of ATP synthesis, ATP hydrolysis-driven proton translocation. And this reaction can be monitored using 9-amino-6-chloro-2-methoxyacridine (ACMA) quenching, a pH-sensitive fluorescent dye (43), as fluorescence decreases along with proton translocation (Fig. 5a). To test this idea, we isolated inverted membrane vesicles from IbsA-expressing cells or vector-expressing cells and measured ATP-mediated proton translocation activity of F_o_F_1_ ATP synthase by fluorescence quenching. The vector-expressing vesicles decreased fluorescence by ∼15% immediately after ATP addition (Fig. 5b). However, IbsA-expressing membrane vesicles resisted the decrease, reducing fluorescence by only ∼5% (Fig. 5b), indicating that IbsA indeed inhibits ATP-driven proton translocation. As controls, vesicles expressing H3A-or L11A-substituted IbsA still showed greater fluorescence quenching than those expressing wild-type IbsA (∼10% and ∼8.5% respectively), but less than the vector control (Fig. 5b). When we used nicotinamide adenine dinucleotide (NADH), an electron carrier, to facilitate proton translocation via the electron transport chain and measured NADH-driven proton translocating activity using the same vesicles, IbsA-expressing vesicles exhibited slightly lower fluorescence quenching than vector-expressing vesicles (Fig. 5c and 5d). However, there was no significant differences between cells expressing wild-type IbsA and IbsA variants (Fig. 5d). This suggests that NADH-driven fluorescence quenching mediated by IbsA expression is not specific to IbsA’s toxin activity or its interaction with F_o_F_1_ ATP synthase.

**Fig. 5.**
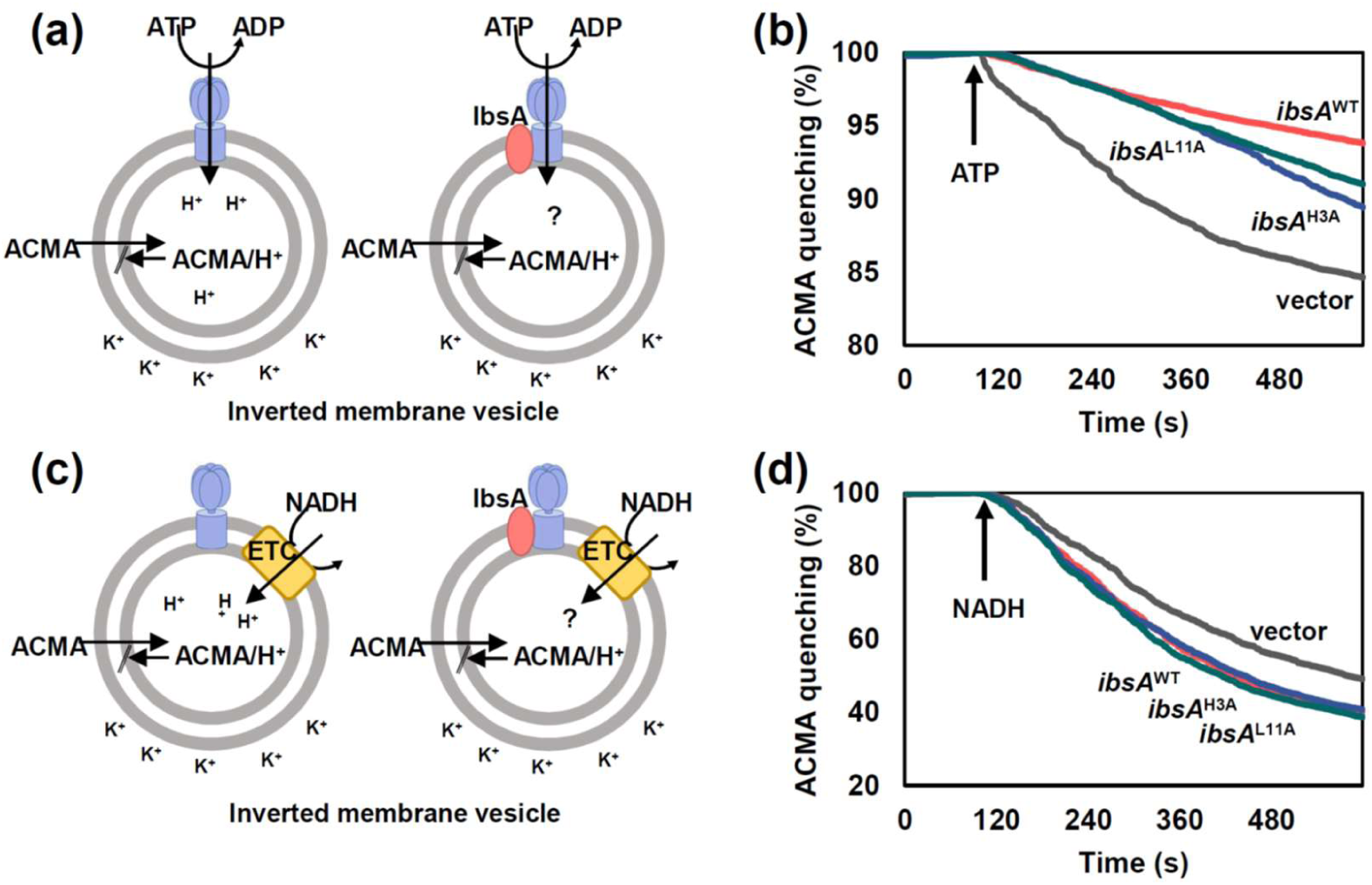
IbsA toxin inhibits ATP-driven proton translocation of F_o_F_1_ ATP synthase (a) Schematic cartoon of the ATP-driven proton translocation assay using ACMA quenching. When the proton is translocated through the F_o_F_1_ ATP synthase in the presence of ATP, ACMA is protonated, leading to fluorescence quenching. (b) Fluorescence quenching of ACMA in inverted membrane vesicles prepared from wild-type harboring either the empty vector or pBAD33-*ibsA*, *ibsA*^H3A^, or *ibsA*^L11A^. The reaction was initiated by adding 1 mM ATP. (c) Schematic cartoon of the NADH-driven proton translocation assay using ACMA quenching. (d) Fluorescence quenching of ACMA in inverted membrane vesicles prepared as in (b), with the reaction initiated by adding 0.5 mM NADH.

To test whether IbsA’s toxin effect is mediated by the F_o_F_1_ ATP synthase, we created a chromosomal *atpF* mutant strain lacking the *b* subunit of F_o_F_1_ ATP synthase. Then, we measured the effect of IbsA expression on growth and ATP levels in the wild-type or *atpF* mutant backgrounds under different carbon sources. In the wild-type background, IbsA expression arrested *Salmonella* growth in succinate-containing media but not in glucose-containing media (Fig. S4a and S4b). This is possibly because IbsA’s growth inhibitory effect can be bypassed when glucose is present, as glucose generates more ATP via both substrate-level phosphorylation and oxidative phosphorylation. By contrast, succinate metabolism is strictly dependent on oxidative phosphorylation, which explains why IbsA’s inhibitory activity was exaggerated in this medium if IbsA functions as a toxin largely dependent on F_o_F_1_ ATP synthase. Next, we tested whether IbsA expression still inhibits growth in the *atpF* mutant background. In contrast to the wild-type, IbsA expression did not exhibited further growth inhibition in the *atpF* mutant both in succinate- or glucose-containing media (Fig. S4c and S4d), indicating that IbsA acts as a toxin largely dependent on the *b* subunit of F_o_F_1_ ATP synthase. Similarly what we detected in the growth phenotype, IbsA expression did not decrease intracellular ATP levels in glucose-, glycerol-, or succinate-containing media when we tested in the *atpF* mutant background (Fig. 3b-3d). Finally, in the inverted vesicles isolated from *atpF* mutant *Salmonella*, IbsA expression did not inhibit ATP-driven proton translocation (Fig. S4e-S4h), supporting that IbsA toxin’s effect is AtpF-dependent.

### *ibsA* RNA levels increase in low Mg^2+^

In the previous section, we showed that IbsA decreases intracellular ATP levels and inhibits bacterial growth (Fig. 2). And, this inhibitory action is mediated by IbsA’s binding to the F_o_ complex of ATP synthase and its inhibition of proton translocation-coupled ATP synthesis (Fig. 4 and 5).

To determine the physiological significance of IbsA expression, we first created a strain lacking the SibA antitoxin RNA. Because SibA antisense RNA overlaps with the *ibsA* gene, we created a chromosomal mutant strain in which both the -10 and -35 promoter sequences of SibA were substituted, potentially eliminating expression of SibA antitoxin RNA and expressing only IbsA toxin (SibA promoter mutant, hereinafter referred to as SibA_pm_)(Fig. 6a). As a control, we also created a strain deleted the DNA sequence of the *ibsA* gene (Δ*ibsA*). Due to the *ibsA*-*sibA* genetic organization, the *ibsA* deletion mutant eliminates the DNA sequences corresponding to both *ibsA* and *sibA* (Fig. 6a).

**Fig. 6.**
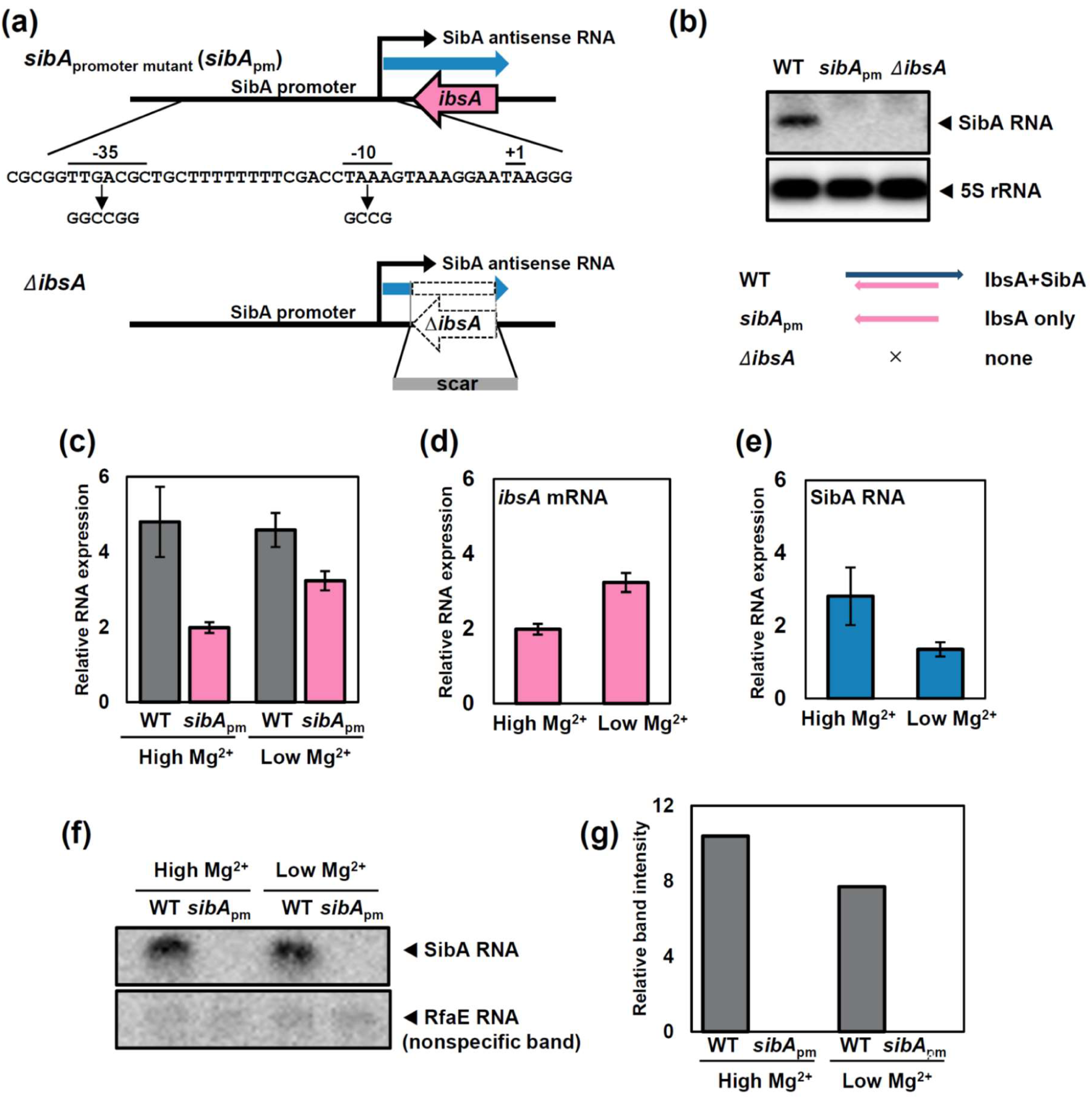
The *sibA* promoter mutant revealed an elevation of *ibsA* mRNA levels in low Mg^2+^ (a) The schematic diagram of the *sibA* promoter mutant and *ibsA* deletion mutant. (b) Northern blot analysis of SibA RNA in wild-type or chromosomal mutant strains with the *sibA* promoter substitution or *ibsA* deletion. RNA samples (10 µg) were analyzed by Northern hybridization using oligonucleotide probes specific to *sibA* and *rrfD* (5s rRNA). Schematic interpretation of the northern result was shown below. (c) Relative levels of SibA RNA isolated from wild-type and the *sibA* promoter mutant strains. Cells were grown at 37°C in N-minimal media containing high (10 mM) or low (0.01 mM) Mg^2+^. Relative RNA levels represent (target RNA/*rrsH* RNA)×10,000. (d, e) *ibsA* (d) and SibA (e) RNA levels calculated from (c). (f) Northern blot analysis of SibA RNA isolated from wild-type and the *sibA* promoter mutant strains. Cells were grown at 37°C in N-minimal media containing high (10 mM) or low (0.01 mM) Mg^2+^. RNA samples (10 ug) were analyzed by Northern hybridization using oligonucleotide probes specific to *sibA*. (g) Relative band intensity was calculated from (f) (SibA band/nonspecific band).

Northern blot analysis showed that the wild-type strain produced SibA, whereas the SibA promoter mutant did not produce SibA RNA, as expected (Fig. 6b). SibA RNA was also undetectable in the *ibsA* mutant, possibly because most of the *sibA* sequence was also deleted (Fig. 6b). As a negative control, 5S rRNA levels were similar across all strains (Fig. 6b).

Given that SYBR green-based RNA quantification detects RNAs from both strands, we measured *ibsA* RNA levels in the wild-type strain (expressing both *ibsA* and SibA RNAs) and in the SibA promoter mutant (expressing *ibsA* RNA only). In low Mg^2+^, an *in vitro* infection-relevant medium, *ibsA* mRNA levels detected in the SibA promoter mutant increased compared to those grown in high Mg^2+^ (Fig. 6c and 6d). Because the wild-type strain exhibited the sum of *ibsA* and SibA RNA levels and did not exhibit significant changes between high and low Mg^2+^ media (Fig. 6c), SibA RNA levels, which were calculated by subtracting *ibsA* RNA levels from total RNA, appeared to decrease in low Mg^2+^ relative to high Mg^2+^ (Fig. 6e). Northern blot analysis using strand-specific probe also showed that the band intensity of SibA RNA slightly decreased in low Mg^2+^ (Fig. 6f and 6g). As a control, *rfaE* RNA readthrough transcripts overlapping with the upstream region of SibA RNA were similar in all tested conditions (Fig. 6f).

### *ibsA* deletion mutant attenuates *Salmonella* virulence in mouse systemic infection

To investigate the function of IbsA toxin during *Salmonella* infection *in vivo*, wild-type, *ibsA* deletion, and *sibA*_pm_ mutant *Salmonella* strains were intraperitoneally injected into mice. Notably, mice infected with the *ibsA* mutant survived significantly longer than the other groups, indicating that *ibsA* deletion attenuates *Salmonella* virulence (Fig. 7a). To explore this further, bacterial burdens in the spleen and liver were examined on days 1, 3, and 6 post-infection. Spleens isolated from wild-type *Salmonella-*infected mice were enlarged compared to PBS-injected mice on days 3 and 6 post-infection, indicating a clear symptom of systemic infection (Fig. 7b). Interestingly, the size and weight of the spleens from *ibsA* mutant-infected mice were similar to those of PBS-injected mice (Fig. 7b and 7c), suggesting that *ibsA* deletion attenuates virulence in this systemic infection model. As a control, spleens isolated from *sibA*_pm_ mutant-infected mice were slightly larger and heavier than those from wild-type-infected mice on days 3 and 6 post-infection (Fig. 7b and 7c). The weight of livers isolated from *sibA*_pm_ mutant-infected mice was slightly higher than in the other three groups (Fig. 7d).

**Fig. 7.**
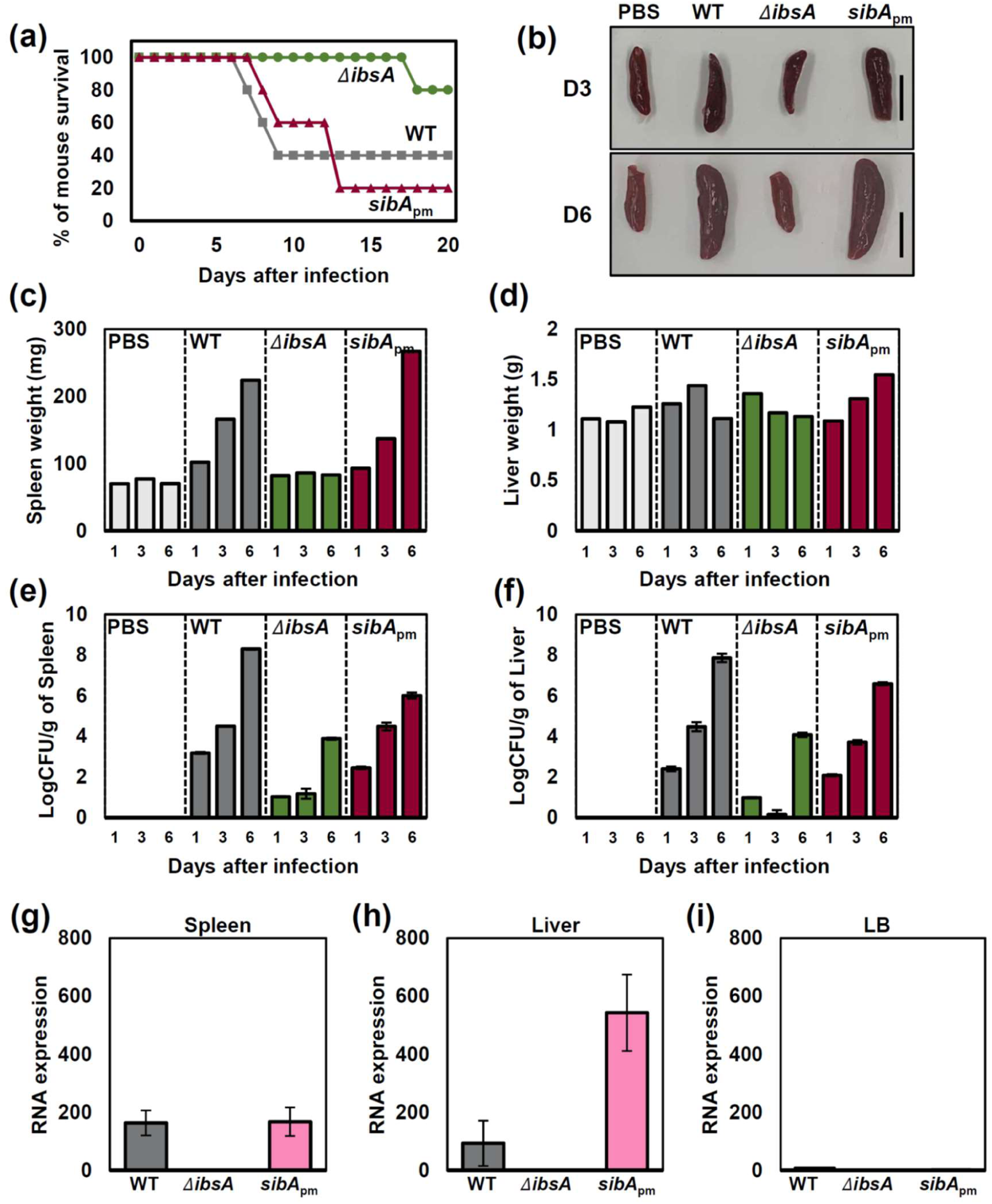
IbsA toxin promotes *Salmonella* virulence in mice (a) Survival of C3H/HeN mice inoculated intraperitoneally with approximately 10^3^ colony-forming units of the wild-type, the *sibA* promoter mutant, or the *ibsA* deletion strains. The data are representative of two independent experiments, which produced similar results. (b) Appearance and relative size of spleens isolated from the mouse groups that received the above *Salmonella* strains or PBS at 3 (D3) or 6 (D6) days post-infection. Shown are representative spleen samples from three independent experiments. Scale bar: 1 cm. (c, d) Differences in organ weights of the mouse groups at 1, 3, and 6 days post-infection. (c) Spleen and (d) liver weights. Control mice received 100 µl of PBS and were negative for *Salmonella* in the spleen and liver. (e, f) Bacterial loads of *Salmonella*-infected spleen and liver. Mouse spleen and liver were harvested at days 1, 3 and 6. Organs were homogenized with PBS containing 0.1% Triton X-100. For measuring the number of bacteria, homogenates were plated on LB solid media with appropriate dilutions. The number of *Salmonella* was determined and represented as Log _10_ CFU/g of organ. (g, h) Relative RNA levels of *Salmonella ibsA-sibA* genes isolated from mouse spleen (g) or liver (h) infected with wild-type, the *sibA* promoter mutant, or the *ibsA* deletion mutant *Salmonella* strains. (i) Relative RNA levels of *Salmonella ibsA-sibA* genes isolated from above strains in LB media. The samples for *in vivo* (spleen and liver) data were obtained at 6 days post-infection. *In vitro* RNA samples (LB) were obtained from *Salmonella* grown to the exponential growth phase (OD_600_ = 0.3) in LB media. RNA expression was determined by measuring (RNA levels of *ibsA* and *sibA* in mouse organ/RNA levels of *rrsH* in mouse organ) ×10,000 or (RNA levels of *ibsA* and *sibA* in LB media/RNA levels of *rrsH* grown in LB media) ×10,000.

Similarly, spleens and livers were homogenized and plated on LB plates to determine colony-forming units (CFU) in each organ. The CFU counts from spleen and liver homogenates of *ibsA* mutant-infected mice were 4 logs lower than those of the wild-type at day 6 post-infection (Fig. 7e and 7f), supporting the notion that *ibsA* deletion decreases bacterial burden at day 6 post-infection. When we measured *ibsA* mRNA levels using the same samples, *ibsA* mRNA levels in spleen and liver homogenates from wild-type *Salmonella*-infected mice were at least 20-fold higher than those from *Salmonella* grown in LB medium (Fig. 7g-7i), indicating that *ibsA* expression is induced in the mouse spleen and liver. It is interesting to note that *ibsA* mRNA levels in the spleen and liver were high relative to SibA RNA, as *ibsA* mRNA levels in the *sibA*_pm_ mutant were similar to or even higher than in the wild-type (Fig. 7g-7i). As a control, *ibsA* mRNA was not detected in *ibsA* mutant-infected mice. These results suggest that IbsA toxin is crucial for *Salmonella* virulence during systemic infection in mice. And this effect is also *atpF*-dependent because *ibsA* deletion did not exhibit a decrease in size or weight of organs, and CFUs when we tested in the *atpF* mutant background (Fig. S5).

### IbsA toxin promotes *Salmonella* transmission during systemic infection

To explore how IbsA toxin affects *Salmonella* systemic infection, we infected mCherry-expressing *Salmonella* into macrophages and measured mean fluorescence intensity to monitor replication efficiency within macrophages (Fig. S6a-S6e). When we measured mCherry fluorescence without macrophage lysis, *ibsA* mutant-infected macrophages exhibited lower fluorescence at 21 h post-infection compared to wild-type-infected macrophages (Fig. 8a). The CFU count of the *ibsA* mutant at 21 h post-infection was also slightly lower than that of wild-type (Fig. 8b top, 8c). As a control, the *sibA*_pm_ mutant exhibited slightly higher mCherry fluorescence than the wild-type at 21 h post-infection (Fig. 8a). However, the formation of nonreplicating cells was not significantly affected by *ibsA* or *sibA* mutations when we measured the percentage of cells expressing high GFP levels in a fluorescence dilution assay (Fig. S6f-S6i).

**Fig. 8.**
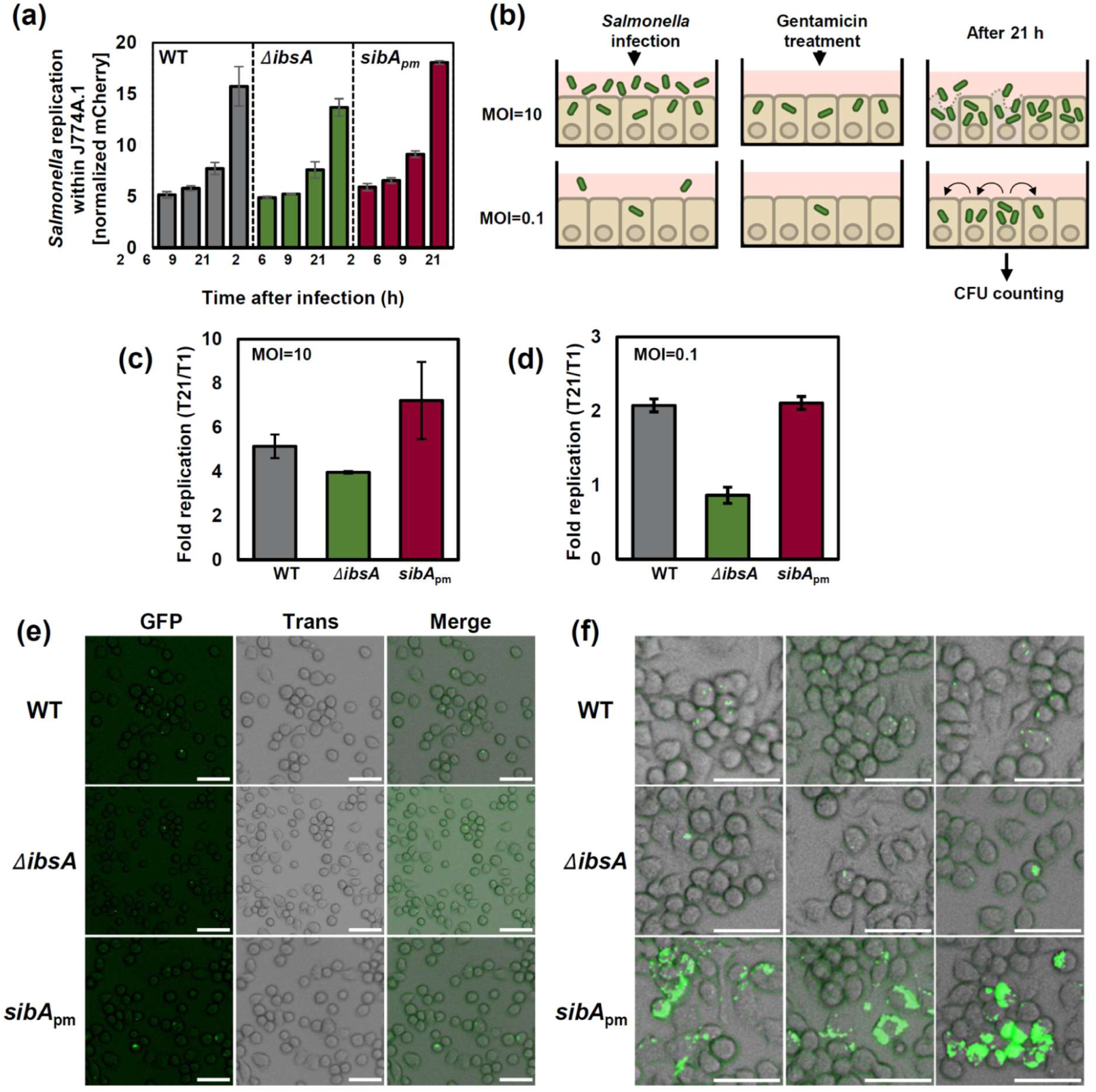
IbsA toxin promotes *Salmonella* transmission during infection (a) Replication inside J774A.1 macrophages of the mCherry-expressing wild-type, *ibsA* deletion, and *sibA* promoter mutant *Salmonella* strains at 2, 6, 9, and 21 h post-infection. *Salmonella* replication was measured as described in Fig. S6a. Normalized mCherry represents [mean fluorescence intensity (MFI) of mCherry-expressing *Salmonella* strains inside macrophages / MFI of mCherry-expressing *Salmonella* strains in overnight-grown LB medium]. See related Fig. S6a-d. (b) Schematic cartoon of *Salmonella* infection at an MOI of 10 (upper) or 0.1 (lower). At a low MOI, *Salmonella* can spread to neighboring macrophages to monitor transmission or multi-round infection (c) Fold replication of single-round infection (MOI of 10) within J774 A.1 macrophage-like cells of wild-type, *ΔibsA,* and *sibA* promoter mutant (*sibA*_pm_) strains at 21 h post-infection. (d) Fold replication of multiple-round infection (MOI of 0.1) within J774 A.1 macrophage-like cells strains listed above at 21 h post infection. Fold replication represents [number of bacteria at T21 / number of bacteria at T1]. Shown are the means and SD from three independent infections. (e) Microscopic images of wild-type, *ΔibsA,* and *sibA*_pm_ *Salmonella* strains expressing GFP inside J774A. 1 cell lines at 2 h post-infection. J774A.1 cells were infected with *Salmonella* strains at a multiplicity of infection (MOI) of 0.1. The cells were incubated for 1 h to allow bacterial invasion, and replication was monitored after additional 1 h incubation. Panels correspond to GFP, Trans (transmitted light), and merged images, respectively. Scale bar: 50 μm. (f) Microscopic images of the same samples at 21 h post-infection. The cells were incubated for 21 h to allow bacterial transmission and were visualized under a fluorescence microscope. Scale bar: 50 μm.

The *ibsA* mutant displayed a slight defect in macrophage survival, but this alone did not fully account for *ibsA*’s defect in the mouse systemic infection model (Fig. 7). We reasoned that high multiplicity of infection (MOI of 10) might not be ideal for examining multiple rounds of infection within macrophages, which mimic *Salmonella*’s transmission to non-infected macrophages (Fig. 8b). Thus, we set up another infection experiment with a lower MOI (MOI of 0.1) to allow *Salmonella* to spread and infect adjacent macrophages (Fig. 8b, bottom). At MOI = 0.1, CFU counts of the wild-type and *sibA* mutant *Salmonella* increased approximately 2-fold at 21 h post-infection (Fig. 8d).

However, the fold replication of the *ibsA* mutant remained close to 1 at 21 post-infection, indicating that its bacterial count of the *ibsA* mutant did not increase during infection (Fig. 8d). To visualize these results, GFP-expressing *Salmonella* was used to infect macrophages and fluorescent images were obtained. Microscopic analysis further supported this notion that GFP signals from the *ibsA* mutant at 21 h post-infection was detected mostly in individual macrophage, whereas GFP signals from wild-type *Salmonella* were detected in multiple adjacent macrophages (Fig. 8e, 8f, S7). These data suggest that *ibsA* deletion has a defect in *Salmonella* transmission during infection. Interestingly, GFP signals from the *sibA*_pm_ mutant *Salmonella* were strongly elevated many adjacent macrophages (Fig. 8f), indicating that increased IbsA expression in the *sibA*_pm_ mutant enhances *Salmonella* spreading between macrophages. This is also in agreement with previous results that the *ibsA* mRNA levels were elevated in the *sibA*_pm_ mutant (Fig. 7g and 7h) and that infected mice exhibited increased organ weight (Fig. 7c and 7d). As a control, at 2 h post-infection, GFP signals were detected as single spots in several individual macrophages, with no significant differences between strains (Fig. 8e).

### IbsA toxin production decreases bacterium’s ATP levels during infection

We then wondered how IbsA toxin promotes *Salmonella* transmission during infection. Given that IbsA interacts with F_o_F_1_ ATP synthase and inhibits proton translocation-coupled ATP synthesis (Fig. 4 and 5), IbsA might enhance *Salmonella* transmission by decreasing bacterium’s ATP levels. To explore this possibility, we first measured IbsA protein levels over time in N-minimal media containing 0.01 mM Mg^2+^, an infection-relevant medium. C-terminally Myc-tagged IbsA proteins were detected at 4 h, slightly higher at 6 h, and maintained at similar levels until 24 h (Fig. 9a), indicating that IbsA toxin proteins are produced under these conditions. Similarly, when we measured IbsA protein levels inside macrophages, IbsA-Myc protein levels were low at 2 h post infection, but elevated at 6, 9, and 21 h post-infection (Fig. 9b). Based on expression kinetics, this suggests that IbsA toxin is required for replication within macrophages or spreading rather than for host cell invasion. As controls, the *ibsA* mutant did not produce IbsA toxin either *in vitro* and *in vivo* conditions (Fig. 9a and 9b).

**Fig. 9.**
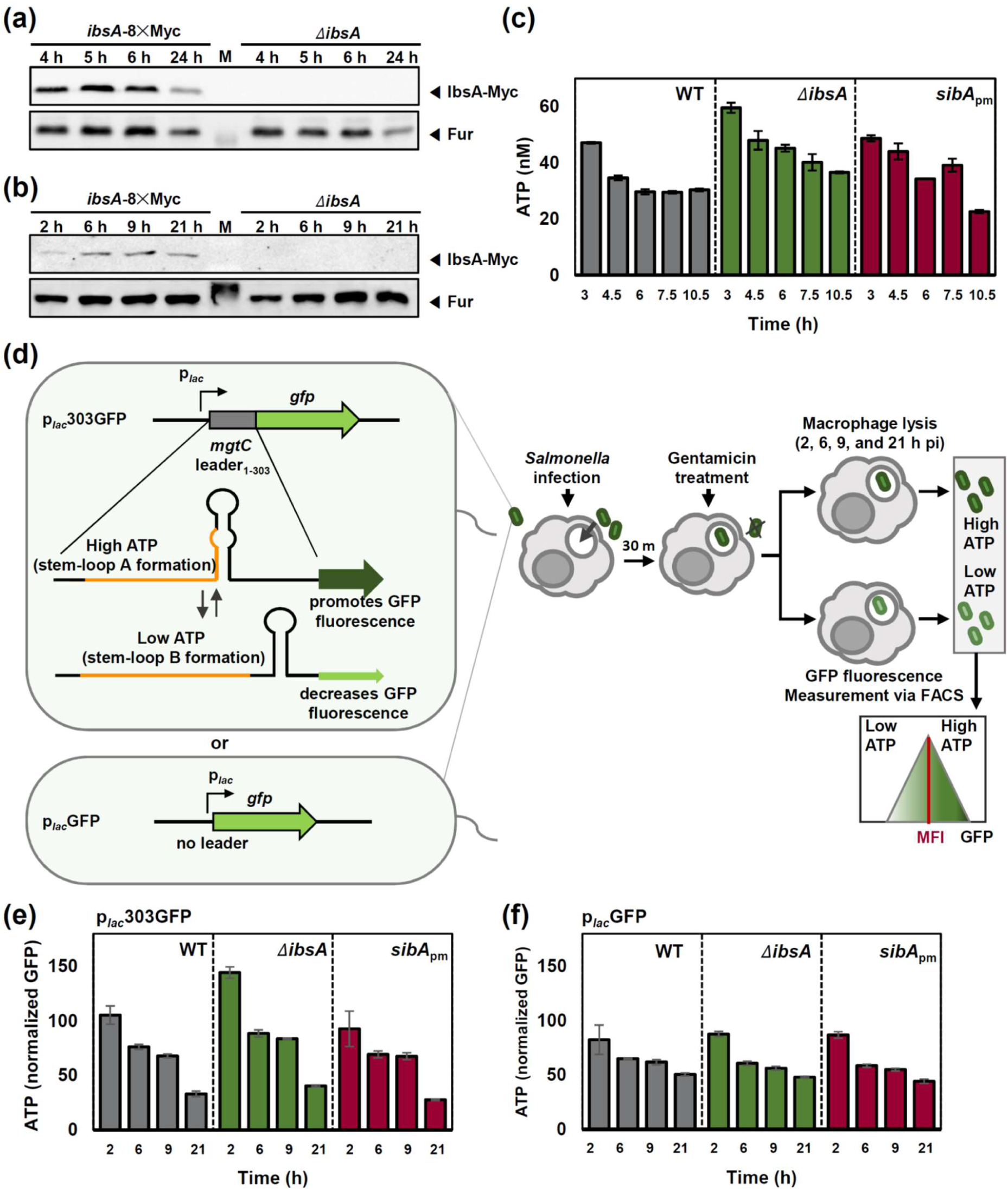
IbsA toxin decreases ATP levels during the course of infection (a) Western blot analysis of crude extracts prepared from *Salmonella* strains with the *ibsA*-8×Myc gene or an *ibsA* deletion. Cells were grown at 37°C in N-minimal medium containing 10 µM Mg^2+^ for the indicated times. Cell lysates were analyzed by immunoblot assay using anti-Myc and anti-Fur antibodies. (b) Western blot analysis of crude extracts prepared from *Salmonella* strains listed in (a) after infection of J774 A.1 macrophage-like cells. Samples were harvested at 2, 6, 9, and 21 h post-infection. Cell lysates were analyzed by immunoblot assay using anti-Myc and anti-Fur antibodies. (c) Temporal changes in intracellular ATP levels of wild-type, *ΔibsA,* and *sibA*_pm_ *Salmonella* strains. The strains were grown at 37°C in N-minimal medium containing 10 µM Mg^2+^ at pH 5.0 (SPI-2 inducing medium). Cells were harvested at the indicated times for measuring ATP levels. (d) Schematic representation of the ATP-responsive p*_lac_*303GFP reporter and measurement of intracellular ATP levels in *Salmonella* harboring this reporter inside macrophages. In this construct, the ATP-responsive leader region of the *mgtC* gene was cloned between a constitutive promoter and the *gfp* gene to monitor bacterial ATP levels inside macrophages. *Salmonella* strains harboring this reporter were used to infect macrophages, and GFP fluorescence was analyzed by flow cytometry to determine mean fluorescence intensity (MFI, red line) at 2, 6, 9, and 21 h post-infection. (e) Temporal changes in intracellular ATP levels of *Salmonella* strains harboring the p*_lac_*303GFP reporter inside J774A.1 macrophages at 2, 6, 9, and 21 h post-infection. Bacterial ATP levels during macrophage infection were analyzed by measuring reporter-driven GFP fluorescence as described in (d). Normalized GFP levels were calculated as [MFI of GFP-expressing *Salmonella* strains inside macrophages / MFI of GFP-expressing *Salmonella* strains grown overnight in LB medium]. See related Fig. S8a-c. (f) Measurement of GFP fluorescence in *Salmonella* strains harboring the p*_lac_*GFP reporter lacking the leader sequence inside macrophages. Infection and GFP analysis were performed as described in (e). See related Fig. S8d-f.

Next, we measured bacterial ATP levels over time. In the infection-relevant medium, wild-type *Salmonella* produced high ATP levels at 3 h, which gradually decreased at 4.5 and 6 h, and then reached a steady state until 10.5 h (Fig. 9c). The *ibsA* mutant accumulated higher ATP levels than the wild-type at 3 h and maintained elevated ATP levels over time, exhibiting a slower decrease in ATP levels (Fig. 9c). By contrast, the *sibA* mutant initially had ATP levels similar to those of wild-type at 3 h, but exhibited a steeper decrease, reaching lower ATP levels than the wild-type at 10.5 h (Fig. 9c). To measure bacterium’s ATP levels inside macrophages, we constructed a plasmid harboring an ATP-responsive leader ahead of the *gfp* gene, which produces GFP fluorescence in response to bacterium’s ATP levels (Fig. 9d) (44). The ATP-responsive GFP reporter was introduced into *Salmonella* strains, which were then used to infect macrophages. Similar to what we detected in the infection-relevant medium, wild-type *Salmonella* inside macrophages exhibited GFP fluorescence at 2 h, which gradually decreased at 6, 9, and 21 h post-infection (Fig. 9e and S8). However, the *ibsA* deletion mutant exhibited higher fluorescence than the wild-type at all time points, indicating that *ibsA* deletion increases bacterium’s ATP levels during infection (Fig. 9e and S8). As controls, the *sibA*_pm_ mutant exhibited slightly lower fluorescence, and a control plasmid lacking the ATP-responsive leader did not exhibit a significant difference in GFP fluorescence between *Salmonella* strains (Fig. 9e, 9f, and S8). These results demonstrate that IbsA toxin expression contributes to maintaining bacterium’s low ATP levels, which could be beneficial for *Salmonella* transmission during systemic infection (Fig. 10).

**Fig. 10.**
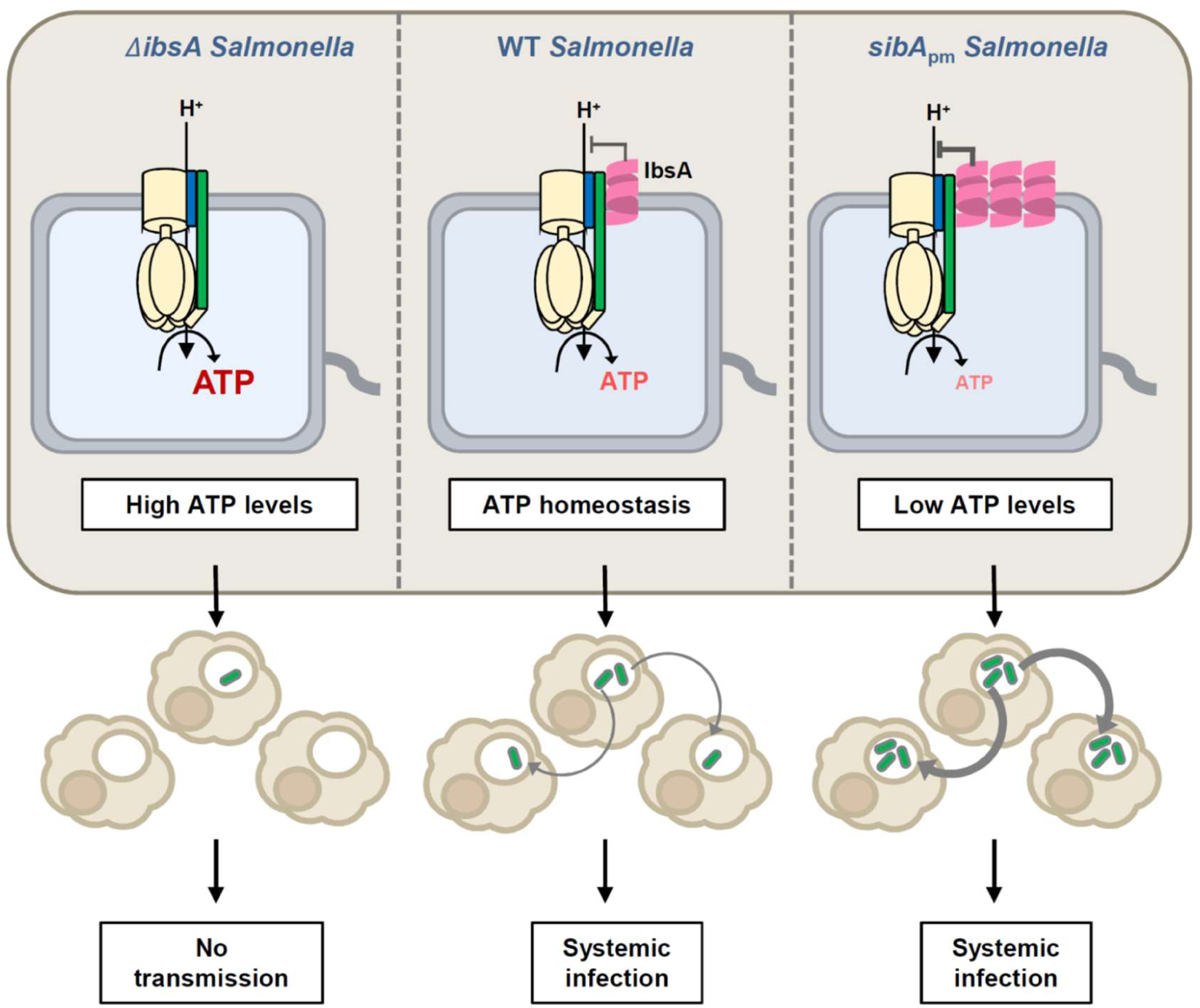
A proposed model for the role of IbsA toxin in *Salmonella* infection The intracellular pathogen *Salmonella* Typhimurium survives and replicates within macrophages, which it could be sensitive to antimicrobials and antibiotics. In the absence of IbsA, the F_o_F_1_ ATP synthase in the bacterial membrane continues to generate high ATP levels using the proton concentration gradient within the macrophage phagosome, which appears to prevent transmission to other macrophages. On the other hand, in the presence of IbsA, the IbsA toxin binds to and inhibits the F_o_ subunit of the F_o_F_1_ ATP synthase by blocking proton translocation, which leads to the maintenance of intracellular pH and ATP levels in the acidic phagosomal compartment. The *sibA*_pm_ mutant lacking SibA antisense RNA produces lower ATP levels, further enhancing *Salmonella* transmission. This IbsA toxin-mediated regulation of bacterium’s ATP levels could reprogram *Salmonella* to transition into a systemic infection.

## Discussion

In this study, we established the function of *Salmonella* IbsA toxin by BACTH screening (Fig. 4). IbsA physically interacts with the F_o_ subunits of F_o_F_1_ ATP synthase (Fig. 4) and inhibits the F_o_F_1_ ATP synthase to translocate protons across the bacterial inner membrane, thereby decreasing ATP synthesis via oxidative phosphorylation (Fig. 5).

The decrease in ATP levels mediated by IbsA toxin contributes to promoting *Salmonella* transmission between macrophages (Fig. 8 and 10) and thus systemic infection (Fig. 7).

In toxin-antitoxin systems, toxins exhibit growth inhibitory effects by interfering with bacterium’s key biological processes when expressed from a plasmid. And, toxin-mediated inhibition of bacterial growth is often associated with persister formation in antibiotic-treated conditions or during host infection (7–9). This is also true for IbsA toxin, given that IbsA overexpression inhibited bacterial growth and led to a ∼10^2^-fold increase in ciprofloxacin-induced persister formation (Fig. 1 and S1). However, the biological function of toxins in their physiological context remains unclear because, in most cases, chromosomal deletion of single or multiple TA systems often does not have a clear phenotype in persister formation (45). This could be due to that the expression of TA loci from its native promoter is low or non-induced in experimental conditions.

In this sense, it is striking that the *ibsA* deletion exhibited a severe defect in mouse systemic infection (Fig. 7), indicating that IbsA actually has a function promoting systemic infection. IbsA’s role seemed to be counterintuitive, considering that most toxins are likely to induce the formation of non-replicating cells or persisters by inhibiting cellular targets. We ascribed this discrepancy between IbsA and other toxins to the following: 1) Although *ibsA* expression from its chromosomal location is induced in low Mg^2+^ media, (Fig. 6), IbsA is unlikely to be expressed at such a high level that it exhibits a *Salmonella* growth inhibitory effect. 2) IbsA toxin is produced inside macrophage at 6, 9, and 21 hours post infection, as well as in the spleen and liver of mice at day 6 post-infection, by which time *Salmonella* is likely to have spread to other macrophages (Fig. 7 and 9).

How does *Salmonella* promote macrophage transmission by decreasing intracellular ATP levels? A clue can be found in a previous report showing that the *Salmonella* virulence factor MgtC increases virulence by repressing ATP levels. *Salmonella* MgtC is highly produced during macrophage infection. Similarly to IbsA, MgtC interacts with the membrane-bound *a* subunit of the F_o_F_1_ ATP synthase and inhibits ATP synthesis (43). Consequently, *Salmonella* strain lacking MgtC continues to accumulate ATP levels, and such an accumulation of ATP levels in turn reroutes metabolic flux toward cellulose production (46). This cellulose production inside macrophages prevents *Salmonella* replication within macrophages by forming biofilms that restrict bacterial motility. Thus, MgtC promotes *Salmonella* replication within macrophages by reducing biofilm production, which could be a similar case for IbsA production inside macrophages.

However, the effect of deletion or production of IbsA appear to exhibit at later stages-during transmission or systemic infection-probably due to differences in expression timing and levels.

HokB, a small membrane toxin in the type I TA system, was previously reported that it oligomerizes and induces pore formation in the membrane, which result in direct ATP leakage through HokB-mediated pores (30). Pore formation and subsequent ATP leakage have been suggested as a common mechanism by which small membrane-associated type I toxins form persisters/non-replicating bacteria, because most of these type I toxins are single membrane-spanning hydrophobic polypeptides ranging from 18 to 52 amino acids in length (24). Although ATP depletion or disruption of the proton motive force appear to be a general mechanism for this type I toxin-mediated persister formation, bacteria can utilize different mechanisms to achieve a non-replicating state. IbsA depletes intracellular ATP levels by inhibiting proton translocation through the F_o_F_1_ ATP synthase, which is coupled to ATP synthesis (Fig. 5). Moreover, bacterial two-hybrid screening identified many potential additional targets that could also result in persister formation (Table 1 and Fig. S3). Therefore, characterization of potential targets could broaden our understanding of the biological functions of type I toxins. And from this IbsA study, it is evident that a toxin’s biological functions must be studied in its biological context.

## MATERIALS AND METHODS

### Bacterial strains, plasmids, oligonucleotides, and growth conditions

Bacterial strains and plasmids used in this study are listed in Table S1. All *S. enterica* serovar Typhimurium strains were derived from the wild-type strain 14028s (47) and were constructed by the one-step gene inactivation method (48) and/or P22-mediated transduction, as previously described (49). DNA oligonucleotides are listed in Table S2. Bacteria were grown at 37°C in Luria-Bertani (LB) broth and N-minimal media (50) supplemented with 0.1% casamino acids, 38 mM glycerol, and the indicated concentrations of MgCl_2_. *E. coli* DH5α was used as the host for plasmid DNA preparation and BTH101 lacking the *cya* gene was used as the host for the bacterial two-hybrid system (51). Ampicillin was used at 50 μg ml^−1^, chloramphenicol at 25 μg ml^− 1^, kanamycin at 50 μg ml^−1^, tetracycline at 10 μg ml^−1^, and fusaric acid (52) at 12 μg ml^−1^. IPTG (isopropyl β-D-1-thiogalactopyranoside) was used at 0.5 mM and L-arabinose at 10 mM.

### Measurement of bacterial growth

Growth of 14028s strains harboring various plasmids was measured. For control, *Salmonella* 14028s containing only vector was used. For growth test, strains were streaked onto solid LB plates and incubated at 37°C. A single colony was then inoculated into 2 ml of LB or N-minimal media containing 10 mM Mg^2+^ with 100 µM chloramphenicol and incubated at 37°C. A 1/100 dilution of pre-cultured samples was inoculated into 15 ml LB or N-minimal media with antibiotics and cultured with shaking at 37°C. When the OD_600_ value reached 0.1, 10 mM L-arabinose was added, and then OD_600_ value was measured every hour. To calculate CFU/ml, cultures were serially diluted every hour and plated onto LB agar. Plates were incubated overnight at 37°C, and CFU were counted.

### Northern blot analysis

To determine the RNA transcript of genes, cells were grown in N-minimal media containing 10 mM Mg^2+^ at 37°C until OD_600_=0.3, and total RNA was isolated using TRIzol^TM^ reagent (Invitrogen) following the manufacturer’s instructions. RNA samples were mixed to an equal volume of RNA sample loading buffer (Sigma-Aldrich, R1386), incubated at 65°C for 5 min, and analyzed by electrophoresis through a formaldehyde-agarose gel in 1× MOPS buffer at 80 V for 1 h. The gel was transferred to a nylon membrane (GE Healthcare, Hybond®-N hybridization membranes, GERPN303N) by capillary action overnight and subjected to UV cross-linking. The transferred membrane was incubated in hybridization buffer (Invitrogen, ULTRAhyb™-Oligo, AM8663) at 42°C for 1 h, followed by the addition of 100 cpm of ^32^P-labeled oligonucleotide probes specific for *myc* or SibA sequences and incubation overnight. The membrane was washed twice for 5 min with low-stringency washing buffer (Invitrogen, NorthernMax™ Low Stringency Wash Buffer, AM8673) and then washed twice for 15 min with preheated high-stringency washing buffer (Invitrogen, NorthernMax™ High Stringency Wash Buffer, AM8674) at 42°C. The membrane was exposed to film for 2 days and visualized using an IP detector (Cytiva, Amersham™ Typhoon Biomolecular Imager, 29187194).

### Western blot analysis

Cells were grown in N-minimal media containing either 10 mM Mg^2+^ or 10 μM Mg^2+^ at 37°C, and crude extracts were prepared in TBS (Tris-buffered saline) buffer by sonication. The samples were electrophoresed on a 12% sodium dodecyl sulfate (SDS)-polyacrylamide gel and transferred onto PVDF membranes. The blots were incubated overnight with monoclonal anti-Myc antibodies (1:5,000 dilution, MBL), anti-GFP antibodies (1:5,000 dilution, Santa Cruz), or polyclonal anti-Fur antibodies (1:10,000 dilution, raised in rabbits) as primary antibodies. The blots were then incubated for 1 h with anti-mouse IgG horseradish peroxidase-linked antibodies (1:10,000 dilution, ThermoFisher) or anti-rabbit IgG horseradish peroxidase-linked antibodies (1:10,000 dilution, ThermoFisher). Protein bands were visualized using the ECL detection system (SuperSignal^®^ West Femto Maximum Sensitivity Substrate, ThermoFisher). The data are representative of at least two independent experiments, which gave similar results.

### Measurement of intracellular ATP

Experiments were performed using the BacTiter-Glo^TM^ Microbial Cell Viability Assay Kit (Promega) following the manufacturer’s instructions with slight modifications. Briefly, cells were grown overnight in N-minimal media containing 10 mM Mg^2+^. A 1/100 dilution of the overnight-grown bacterial culture was inoculated in 15 ml of N-minimal media containing 10 mM Mg^2+^ and grown for 3 h. Then, 10 mM L-arabinose was added and the culture was grown for an additional 2 h. Cells were normalized by measuring OD_600_ and resuspended in 1 ml of PBS (phosphate-buffered saline). Next, 80 μl of this cell suspension was dispensed into an opaque 96-well microplate (SPL), followed by the addition of 80 μl of BacTiter-Glo^TM^ reagent. The contents were briefly mixed by pipetting and incubated for 5 min. The luminescence of the samples was measured using a Synergy H1 plate reader (BioTek).

### Measurement of membrane potential

Experiments were performed using the BacLight^TM^ Bacterial Membrane Potential Kit (Invitrogen) following the manufacturer’s instructions with slight modifications. Briefly, cells were grown similarly to those for measuring intracellular ATP. For measuring membrane potential, 150 μl of the normalized cell suspension was aliquoted into a 96-well black/clear-bottom plate (ThermoFisher), followed by the addition of 1.5 μl of 3 mM DiOC_2_ (3,3’-diethyloxacarbocyanine Iodide) in the presence or absence of 1.5 μl of 500 μM of CCCP (carbonyl cyanide 3-chlorophenylhydrazon). The contents were briefly mixed by pipetting and incubated for 30 min at 37°C. Membrane potential was determined by measuring the absorbance of the solutions at 645 nm and 530 nm using a Synergy H1 plate reader (BioTek).

### Bacterial two-hybrid assay

To screen for target proteins that interact with the IbsA toxin *in vivo*, a bacterial two-hybrid (BACTH) assay was conducted as described (51). The *Salmonella* random genomic DNA (gDNA) fragment library was constructed using two different methods. First, gDNA samples digested with the Sau3AI restriction enzyme for 15, 30, and 45 min were cloned into pUT18 or pUT18c plasmids digested with BamHI. Second, gDNA samples fragmented by sonication for 2, 4, and 6 min were blunted using the Klenow fragment of DNA polymerase I (NEB) and cloned into pUT18 or pUT18c plasmids digested with SmaI. The *Escherichia coli* BTH101 strain lacking the *cya* adenylate cyclase gene (-*cya*) was co-transformed with derivatives of pUT18 or pUT18c carrying random DNA fragments and pKT25 carrying *ibsA*. Transformants were plated on MacConkey agar containing 100 μM IPTG, 100 μM ampicillin, and 100 μM kanamycin, followed by incubation at 30°C for 48 h. Red colonies were re-streaked on the same MacConkey plates, and the genomic DNA fragments in the pUT18 or pUT18c plasmids were verified by DNA sequencing. To validate IbsA-interacting candidates, a strain harboring the pKT25-MgtR peptide and the pUT18-MgtC protein, which were identified as interacting membrane proteins (53, 54), was used as a positive control.

### Immunoprecipitation assay

The interaction between the IbsA protein and the ATP synthase F_o_ subunit was investigated in wild-type *Salmonella* expressing the *ibsA* gene derivatives with N-terminally GFP-tagged constructs under an arabinose-inducible promoter (pBAD33-GFP-*ibsA*, pBAD33-GFP-*ibsA*^His3Ala^, or pBAD33-GFP-*ibsA*^Leu11Ala^) and C-terminally HA-tagged *atpF* gene from its chromosomal locus. Cells were grown overnight in LB media. A 1/100 dilution of the overnight grown bacterial culture was inoculated in 15 ml of LB media and grown for 1 h. Then, 1 mM L-arabinose added and the culture was grown for an additional 2 h. Cells were normalized by measuring OD_600_. Crude extracts were prepared in TBS (Tris-buffered saline) buffer by sonication. For a pull-down assay with anti-HA antibodies, 50 μl of the crude extracts were kept as input and 350 μl of the protein extracts were mixed with 25 μl of EZview^TM^ Red anti-HA Affinity Gel (Sigma-Aldrich) overnight at 4°C on a nutator (BenchMark). Beads were washed five times with TBS washing buffer, and the bound proteins were eluted in SDS sample buffer. The eluates were resolved on 12% SDS-polyacrylamide gels, transferred to a nitrocellulose membrane, and analyzed by Western blot using anti-HA (1:5,000 dilution, Rockland) and anti-GFP (1:5,000 dilution, Santa Cruz) antibodies overnight. The blots were developed by incubation with anti-mouse IgG horseradish peroxidase-linked antibodies (1:10,000 dilution, ThermoFisher) or anti-rabbit IgG horseradish peroxidase-linked antibodies (1:10,000 dilution, ThermoFisher) for 1 h and were detected using the ECL detection system (SuperSignal® West Femto Maximum Sensitivity Substrate, ThermoFisher).

### Inverted membrane vesicle preparation

Cells were grown overnight in N-minimal medium containing 10 mM Mg^2+^. A 1/100 dilution of the overnight-grown bacterial culture was inoculated into 15 ml of N-minimal medium containing 10 mM Mg^2+^ and grown for 3 h. Then, 10 mM L-arabinose was added and the culture was grown for an additional 2 h. Cells were normalized by measuring OD_600_. Crude extracts were prepared in lysis buffer (10 mM HEPES/KOH, pH 7.5, 5 mM MgCl_2_, 10% glycerol) and disrupted by sonication. Supernatants were centrifuged for 2 h at 240,000 × *g* (Optima XE-90 Ultracentrifuge, Type SW 55 Ti Rotor, Beckman Coulter). The pellets were resuspended in the same lysis buffer. The protein concentration in the prepared membrane fractions was determined using a NanoDrop spectrophotometer (ThermoFisher).

### Measurement of proton translocating activity of the F_1_F_o_ ATP synthase

ATP-driven proton translocating activity was determined by monitoring fluorescence quenching of ACMA (9-amino-6-chloro-2-methoxyacridine), a pH-dependent dye, as described with modifications (Suzuki et al., 2007)(43). Membrane vesicles were diluted to a concentration of 30 μg of protein per ml of assay buffer (10 mM HEPES/ KOH, pH7.5, 100 mM KCl, 5 mM MgCl_2_, and 0.3 µg/ ml of ACMA). The proton translocation reaction was initiated by adding 1 mM ATP or 0.5 mM NADH, and fluorescence quenching was monitored at room temperature with excitation at 410 nm and emission at 490 nm using an LS-55 fluorescence spectrometer (PerkinElmer).

### Measurement of *ibsA* transcript levels by real-time polymerase chain reaction

Total RNA was isolated using the RNeasy Kit (Qiagen) according to the manufacturer’s instructions. The purified RNA was quantified using a NanoDrop spectrophotometer (NanoDrop Technologies). Complementary DNA (cDNA) was synthesized using the PrimeScriptTM RT Reagent Kit (TaKaRa). The mRNA levels of the *ibsA* gene were measured by quantifying the cDNA using SYBR Green PCR Master Mix (TOYOBO) and the appropriate primers and monitored using the StepOnePlus Real-Time PCR System (Applied Biosystems). The mRNA levels of the target gene were calculated using a standard curve generated from *Salmonella* 14028s genomic DNA, and the data were normalized to the levels of 16S ribosomal RNA.

### Mouse virulence assay

Six-to eight-week-old female C3H/HeN mice were inoculated intraperitoneally with ∼10^3^ colony forming units of *Salmonella* strains. Mouse survival was monitored for 21 days. Virulence assays were conducted twice with similar outcomes and the data correspond to groups of five mice. To examine bacterial burden in mice, infected mice were sacrificed at the indicated time points post-infection and the spleens and livers were dissected, homogenized in PBS buffer containing 0.1% Triton X-100. Each homogenate was properly diluted, plated on LB agar plates, and incubated overnight at 37°C. All animals were housed in a temperature- and humidity-controlled room, with a 12-h light/12-h dark cycle. The experimental procedures were approved by the Korea University Institutional Animal Care and Use Committee.

### Measuring gene expression inside mouse

The spleen and livers were harvested from *Salmonella*-infected mice, lysed, and stabilized with TRIzol^TM^ reagent (Invitrogen™). RNA was prepared according to manufacturer’s instructions. Control RNA was obtained from *Salmonella* grown to an exponential growth phase (OD_600_=0.3) in LB medium. RNA expression *in vivo*/*in vitro* was determined by measuring (mRNA levels of the gene inside organs/mRNA levels of *rrsH* inside organs) / (mRNA levels of the gene in LB medium/mRNA levels of *rrsH* grown in LB medium).

### Flow cytometry

Macrophage infection and assessment of *Salmonella* replication were performed using the macrophage-like cell line J774.A1. J774.A1 macrophages were grown in Dulbecco’s Modified Eagle Medium (DMEM) supplemented with 10% (vol/vol) fetal bovine serum (FBS) at 37°C and 5% (vol/vol) CO_2_ in a T75 flask. Prior to infection, 5×10^5^ macrophages were seeded in 24-well plates and incubated for 20 h. *Salmonella* carrying pFCcGi, an mCherry-constitutive and GFP-inducible plasmid, or p*_lac_*303GFP, a plasmid with the *gfp* gene expressed from a constitutive promoter and a 303-nt-long leader sequence from the *mgtCBRU* operon (44), were grown overnight in LB medium and used to infect macrophages at a multiplicity of infection (MOI) of 10. The plates were centrifuged at 1500 rpm for 5 min at room temperature and incubated for an additional 30 min. Then, the extracellular bacteria were washed three times with PBS (phosphate-buffered saline) and killed by incubation with DMEM supplemented with 10% FBS and 120 μg/ml gentamicin for 1 h. After 1h, the DMEM was replaced with fresh DMEM containing 12 μg/ml gentamicin, and the incubation was continued at 37°C. At the indicated time points, infected macrophages were washed and detached using a cell scrapper with PBS. The fluorescence of bacteria inside macrophages was subsequently assessed by FACS analysis. Samples were analyzed on a NovoCyte^TM^ Flow Cytometer (ACEA) using NovoExpress® software (ACEA). On the NovoCyte^TM^ Flow Cytometer, fluorophores were excited at a wavelength of 488 nm, with red fluorescence detected at 615 nm and green fluorescence at 530 nm. Data were analyzed with NovoExpress® software.

### Macrophage survival and transmission assay

Intramacrophage survival and transmission assays were performed with the following modifications (43). 5×10^5^ J774 A.1 macrophages were seeded in 24-well plates and cultured at 37°C. Overnight-grown bacteria were added to the macrophages at an MOI of 10 for the intracellular survival assay and an MOI of 0.1 for the transmission assay. To measure the number of bacteria at the indicated time points, cells were lysed with PBS containing 0.1% Triton X-100 and plated on LB plates with appropriate dilutions. The percentage survival or transmission was obtained by dividing the number of bacteria recovered after 21 h by the number of bacteria recovered at 1 h. All experiments were performed in triplicate and the results are representative of three independent experiments.

### Fluorescent image analysis of *Salmonella* infection

To obtain fluorescent images of *Salmonella* infection or transmission, 1×10^6^ J774. A1 macrophages were seeded in 6-well plates. For bacterial culture, *S.* Typhimurium strains harboring the pFPV25.1 vector were inoculated into 2 ml of LB broth with 100 μM ampicillin and incubated at 37°C overnight. Overnight-grown bacteria were added to the macrophages at an MOI of 0.1. Fluorescent images were obtained using the EVOS FL AUTO imaging system (Thermo Scientific).

### Construction of a strain with the chromosomal deletion

*Salmonella* strains deleted for the *ibsA* or *atpF* genes were generated by the one-step gene inactivation method (48). The Km^R^ cassettes for the *ibsA* and *atpF* genes were PCR-amplified from plasmid pKD4 using primers SZ008/SZ009 (for *ibsA*) and SZ100/SZ099 (for *atpF*). The resulting PCR products were integrated into the 14028s chromosome to generate SY050 (*ibsA*::Km^R^) and SY199 (*ibsA*::Km^R^). The *ibsA* strain (SM051) and *atpF* strain (SY202) were generated by removing the Km^R^ cassettes from SY050 and SY199 via plasmid pCP20 as described (48).

### Construction of strains with chromosomal tagged genes

A *Salmonella* strain with an 8×Myc-tagged *ibsA* and a 2×HA-tagged *atpF* gene were generated as previously described (55, 56). The Km^R^ cassettes of pBOP508 (8×Myc) and pKD13-2×HA (2×HA) were amplified by PCR using SZ173/SZ174 (for *ibsA*-8×Myc) and SZ098/SZ099 (for *atpF*-2×HA). The subsequent PCR products were integrated into wild-type 14028s harboring pKD46 to generate SY325 (*ibsA*-8×Myc::Km^R^) and SY200 (*atpF*-2×HA::Km^R^). The *ibsA*-8×Myc (SY329) and *atpF*-2×HA (SY203) strains were constructed by removing the Km^R^ cassettes using the pCP20 plasmid.

### Construction of strains with chromosomal substitutions in the *ibsA* gene and *sibA* promoter

To generate strains with chromosomal mutations in the *ibsA* coding region and *sibA* promoter, we used the fusaric acid-based counter selection method (57). First, we introduced Tet^R^ cassettes into the *ibsA/sibA* regions as follow: we generated PCR products harboring the *tetRA* gene using primers SZ046/SZ047 and MS7953s chromosomal DNA as a template. The purified PCR product was used to electroporate the 14028s chromosome containing plasmid pKD46 (48). The resulting *ibsA/sibA*::*tetRA* (SY100) strain containing plasmid pKD46 was kept at 30°C. Then, we prepared DNA fragments carrying various nucleotide substitutions in the *ibsA* gene by a two-step PCR process to replace the *tetRA* cassettes. In the first PCR reaction, the primer pairs SZ111/SZ108 and SZ107/SZ112 (for the 3^rd^ histidine codon to alanine), SZ111/SZ110 and SZ109/SZ112 (for the 11^th^ leucine codon to alanine), and SZ111/SZ059 and SZ058/SZ112 (for the *sibA* promoter mutation) were used with 14028s genomic DNA as a template. In the second PCR reaction, we mixed the two PCR products from the first PCR reaction as templates and amplified DNA fragments using primers SZ111 and SZ112. The resulting PCR products were purified and integrated into the SY100 (*ibsA/sibA*::*tetRA*) chromosome harboring pKD46. Selection against Tet^R^ with media containing fusaric acid generated SY246 (*ibsA* ^His 3 Ala^), SY247 (*ibsA* ^Leu 11 Ala^), and SY168 (*sibA*), tetracycline-sensitive, ampicillin-sensitive (Tet^S^ Amp^S^) chromosomal mutants. The expected nucleotide substitutions were verified by DNA sequencing.

### Plasmid construction

To determine the start codon of *ibsA*, the plasmids pBAD33-*ibsA*^GTG-started^-8×Myc, pBAD33-*ibsA*^ATG-started^-8×Myc, pBAD33-*ibsA*^GTG>TAG^, and pBAD33-*ibsA*^ATG>GCG^ were constructed as follows: DNA fragments corresponding to *ibsA*-8×Myc genes were amplified by PCR using the primer pairs, SZ176/SZ178 (for ibsA^GTG-started^-8×Myc) and SZ177/SZ178 (for ibsA^ATG-started^-8×Myc), using SY329 (*ibsA*-8×Myc) as a template. DNA fragments carrying nucleotide substitutions in the *ibsA* gene (*ibsA*^ATG>GCG^) were generated by a two-step PCR process. For the first PCR reaction, the primer pairs SZ001/SZ014 and SZ013/SZ002 were used. For the second PCR reaction, SZ001/SZ002 (for *ibsA*^ATG>GCG^) were used with the first PCR products as templates. A DNA fragment carrying *ibsA*^GTG>TAG^ substitution was amplified using the primer pairs SZ012/SZ002 (for *ibsA*^GTG>TAG^) using 14028s genomic DNA as a template. After purification, the PCR products were digested with KpnI and HindIII and cloned into pBAD33 plasmids digested with the same enzymes.

For alanine scanning, the plasmids pBAD33-*ibsA*, pBAD33-*ibsA* ^Met 2 Ala^, pBAD33-*ibsA* ^His 3 Ala^, pBAD33-*ibsA* ^Gln 4 Ala^, pBAD33-*ibsA* ^Val 5 Ala^, pBAD33-*ibsA* ^Ile 6 Ala^, pBAD33-*ibsA* ^Ile 7 Ala^, pBAD33-*ibsA* ^Leu 8 Ala^, pBAD33-*ibsA* ^Ile 9 Ala^, pBAD33-*ibsA* ^Val 10 Ala^, pBAD33-*ibsA* ^Leu 11 Ala^, pBAD33-*ibsA* ^Leu 12 Ala^, pBAD33-*ibsA* ^Leu 13 Ala^, pBAD33-*ibsA* ^Ile 14 Ala^, pBAD33-*ibsA* ^Ser 15 Ala^, pBAD33-*ibsA* ^Phe 16 Ala^, and pBAD33-*ibsA* ^Tyr 19 Ala^ were constructed as follows: DNA fragments corresponding to *ibsA* and its variants were amplified by PCR using specific primer pairs (SZ003/SZ002 (for *ibsA*), SZ019/SZ002 (for *ibsA* ^Met 2 Ala^), SZ020/SZ002 (for *ibsA* ^His 3 Ala^), SZ021/SZ002 (for *ibsA* ^Gln 4 Ala^), SZ022/SZ002 (for *ibsA* ^Val 5 Ala^), SZ023/SZ002 (for *ibsA* ^Ile 6 Ala^), SZ024/SZ002 (for *ibsA* ^Ile 7 Ala^), SZ025/SZ002 (for *ibsA* ^Leu 8 Ala^), SZ026/SZ002 (for *ibsA* ^Ile 9 Ala^), SZ027/SZ002 (for *ibsA* ^Val 10 Ala^), SZ003/SZ028 (for *ibsA* ^Leu 11 Ala^), SZ003/SZ029 (for *ibsA* ^Leu 12 Ala^), SZ003/SZ030 (for *ibsA* ^Leu 13 Ala^), SZ003/SZ031 (for *ibsA* ^Ile 14 Ala^), SZ003/SZ032 (for *ibsA* ^Ser 15 Ala^), SZ003/SZ033 (for *ibsA* ^Phe 16 Ala^), and SZ003/SZ034 (for *ibsA* ^Tyr 19 Ala^)), using 14028s genomic DNA as a template. After purification, the PCR products were digested with KpnI and HindIII and cloned into pBAD33 plasmids digested with the same enzymes.

For the bacterial two-hybrid assay, the plasmids pKT25-*ibsA,* pKT25-*ibsA* ^His 3 Ala^, pKT25-*ibsA* ^Leu 11 Ala^, and pUT18-*atpF* were constructed as follows: DNA fragments corresponding to the *ibsA, ibsA* ^His 3 Ala^*, ibsA* ^Leu 11 Ala^, and *atpF* genes were amplified by PCR using the primer pairs SZ007/SZ006 (for *ibsA*), SZ070/SZ006 (for *ibsA* ^His 3 Ala^), SZ007/SZ006 (for *ibsA* ^Leu 11 Ala^), and SZ037/SZ038 (for *atpF*), using 14028s (for *ibsA* and *atpF*), SY246 (for *ibsA* ^His 3 Ala^), and SY247 (for *ibsA* ^Leu 11 Ala^) genomic DNAs as templates, respectively. The amplified DNA fragments were digested with XbaI and KpnI and cloned into pKT25 or pUT18, which were digested with the same enzymes. The sequences of the resulting constructs were verified by DNA sequencing.

To generate an N-terminally GFP-tagged construct, the plasmid pBAD33-GFP (pEN203) was constructed as follows: A DNA fragment corresponding to the *gfp* gene lacking the stop codon, was amplified by PCR using the primer pair KU234 / KU238 with the plasmid pFPV25 as a template. After purification, the PCR product was digested with KpnI and XbaI and cloned into the pBAD33 plasmid digested with the same enzymes.

For the immunoprecipitation assay, the plasmids pBAD33-GFP-*ibsA*, pBAD33-GFP-*ibsA* ^His 3 Ala^, and pBAD33-GFP-*ibsA* ^Leu 11 Ala^ were constructed as follows: DNA fragments corresponding to *ibsA* and its variants were amplified by PCR using the primer pairs SZ068/SZ002 (for *ibsA*), SZ105/SZ002 (for *ibsA* ^His 3 Ala^), and SZ068/SZ002 (for *ibsA* ^Leu 11 Ala^), using 14028s (for *ibsA*), SY246 (for *ibsA* ^His 3 Ala^), and SY247 (for *ibsA* ^Leu 11 Ala^) genomic DNAs as templates, respectively. The amplified DNA fragments were digested with XbaI and HindIII and cloned into pBAD33-GFP digested with the same enzymes. The sequences of the resulting constructs were verified by DNA sequencing.

To measure intracellular ATP levels during mouse infection, p*_lac_*303GFP, a plasmid containing the constitutive p*_lac1-6_* promoter and the 303 nt-long leader region of the *mgtC* gene fused to a promoterless *gfp* gene, was constructed as follows: A PCR fragment was generated with primers 9870 and 8117 using 14028s genomic DNA as a template. The DNA fragment was then digested with EcoRI and XbaI and cloned into the pfpV25 plasmid digested with the same enzymes.

## DATA AVAILABILITY

All other relevant data are available from the corresponding author upon reasonable request.

## ACKNOWLEDGMENTS

This work was supported by Basic Science Research Program through the National Research Foundation of Korea (NRF) funded by the Ministry of Science, ICT and Future Planning [NRF-2022R1A2B5B02002256 and NRF-2022R1A4A1025913 to E.-J.L.] and a grant from Korea University.

## Author contributions

Conceptualization: E.-J.L.

Investigation: SY.K. SG.K.

Supervision: E.-J.L.

Writing—original draft: E.-J.L. SY.K.

Writing—review & editing: E.-J.L.

## CONFLICT OF INTEREST STATEMENT

The authors declare no conflict of interest.

## Notes

### Competing Interest Statement

The authors have declared no competing interest.

## REFERENCES

1. Steele-Mortimer O, Méresse S, Gorvel JP, Toh BH, Finlay BB. 1999. Biogenesis of Salmonella typhimurium-containing vacuoles in epithelial cells involves interactions with the early endocytic pathway. Cell Microbiol 1:33–49.

2. Ohl ME, Miller SI. 2001. Salmonella: a model for bacterial pathogenesis. Annu Rev Med 52:259–74.

3. Kumar Y, Valdivia RH. 2009. Leading a sheltered life: intracellular pathogens and maintenance of vacuolar compartments. Cell Host Microbe 5:593–601.

4. Pradhan D, Devi Negi V. 2019. Stress-induced adaptations in Salmonella: A ground for shaping its pathogenesis. Microbiol Res 229:126311.

5. Helaine S, Cheverton AM, Watson KG, Faure LM, Matthews SA, Holden DW. 2014. Internalization of Salmonella by macrophages induces formation of nonreplicating persisters. Science 343:204–8.

6. Wiradiputra MRD, Khuntayaporn P, Thirapanmethee K, Chomnawang MT. 2022. Toxin-Antitoxin Systems: A Key Role on Persister Formation in Salmonella enterica Serovar Typhimurium. Infect Drug Resist 15:5813–5829.

7. Gollan B, Grabe G, Michaux C, Helaine S. 2019. Bacterial Persisters and Infection: Past, Present, and Progressing. Annu Rev Microbiol 73:359–385.

8. Harms A, Brodersen DE, Mitarai N, Gerdes K. 2018. Toxins, Targets, and Triggers: An Overview of Toxin-Antitoxin Biology. Mol Cell 70:768–784.

9. Wilmaerts D, Windels EM, Verstraeten N, Michiels J. 2019. General Mechanisms Leading to Persister Formation and Awakening. Trends Genet 35:401–411.

10. Gerdes K, Bech FW, Jørgensen ST, Løbner-Olesen A, Rasmussen PB, Atlung T, Boe L, Karlstrom O, Molin S, von Meyenburg K. 1986. Mechanism of postsegregational killing by the hok gene product of the parB system of plasmid R1 and its homology with the relF gene product of the E. coli relB operon. Embo j 5:2023–9.

11. Gerdes K, Rasmussen PB, Molin S. 1986. Unique type of plasmid maintenance function: postsegregational killing of plasmid-free cells. Proc Natl Acad Sci U S A 83:3116–20.

12. Gerdes K, Maisonneuve E. 2012. Bacterial persistence and toxin-antitoxin loci. Annu Rev Microbiol 66:103–23.

13. Yamaguchi Y, Park JH, Inouye M. 2011. Toxin-antitoxin systems in bacteria and archaea. Annu Rev Genet 45:61–79.

14. Page R, Peti W. 2016. Toxin-antitoxin systems in bacterial growth arrest and persistence. Nat Chem Biol 12:208–14.

15. Brantl S, Jahn N. 2015. sRNAs in bacterial type I and type III toxin-antitoxin systems. FEMS Microbiol Rev 39:413–27.

16. Fozo EM, Hemm MR, Storz G. 2008. Small toxic proteins and the antisense RNAs that repress them. Microbiol Mol Biol Rev 72:579–89, Table of Contents.

17. Berghoff BA, Wagner EGH. 2017. RNA-based regulation in type I toxin-antitoxin systems and its implication for bacterial persistence. Curr Genet 63:1011–1016.

18. Kawano M, Aravind L, Storz G. 2007. An antisense RNA controls synthesis of an SOS-induced toxin evolved from an antitoxin. Mol Microbiol 64:738–54.

19. Darfeuille F, Unoson C, Vogel J, Wagner EG. 2007. An antisense RNA inhibits translation by competing with standby ribosomes. Mol Cell 26:381–92.

20. Hayes F. 2003. Toxins-antitoxins: plasmid maintenance, programmed cell death, and cell cycle arrest. Science 301:1496–9.

21. Masachis S, Darfeuille F. 2018. Type I Toxin-Antitoxin Systems: Regulating Toxin Expression via Shine-Dalgarno Sequence Sequestration and Small RNA Binding. Microbiol Spectr 6.

22. Weel-Sneve R, Kristiansen KI, Odsbu I, Dalhus B, Booth J, Rognes T, Skarstad K, Bjørås M. 2013. Single transmembrane peptide DinQ modulates membrane-dependent activities. PLoS Genet 9:e1003260.

23. Fozo EM, Makarova KS, Shabalina SA, Yutin N, Koonin EV, Storz G. 2010. Abundance of type I toxin-antitoxin systems in bacteria: searches for new candidates and discovery of novel families. Nucleic Acids Res 38:3743–59.

24. Shore SFH, Leinberger FH, Fozo EM, Berghoff BA. 2024. Type I toxin-antitoxin systems in bacteria: from regulation to biological functions. EcoSal Plus doi:10.1128/ecosalplus.esp-0025-2022:eesp00252022.

25. Guo Y, Quiroga C, Chen Q, McAnulty MJ, Benedik MJ, Wood TK, Wang X. 2014. RalR (a DNase) and RalA (a small RNA) form a type I toxin-antitoxin system in Escherichia coli. Nucleic Acids Res 42:6448–62.

26. Verstraeten N, Knapen WJ, Kint CI, Liebens V, Van den Bergh B, Dewachter L, Michiels JE, Fu Q, David CC, Fierro AC, Marchal K, Beirlant J, Versées W, Hofkens J, Jansen M, Fauvart M, Michiels J. 2015. Obg and Membrane Depolarization Are Part of a Microbial Bet-Hedging Strategy that Leads to Antibiotic Tolerance. Mol Cell 59:9–21.

27. Unoson C, Wagner EG. 2008. A small SOS-induced toxin is targeted against the inner membrane in Escherichia coli. Mol Microbiol 70:258–70.

28. Yamaguchi Y, Tokunaga N, Inouye M, Phadtare S. 2014. Characterization of LdrA (long direct repeat A) protein of Escherichia coli. J Mol Microbiol Biotechnol 24:91–7.

29. Fozo EM, Kawano M, Fontaine F, Kaya Y, Mendieta KS, Jones KL, Ocampo A, Rudd KE, Storz G. 2008. Repression of small toxic protein synthesis by the Sib and OhsC small RNAs. Mol Microbiol 70:1076–93.

30. Wilmaerts D, Bayoumi M, Dewachter L, Knapen W, Mika JT, Hofkens J, Dedecker P, Maglia G, Verstraeten N, Michiels J. 2018. The Persistence-Inducing Toxin HokB Forms Dynamic Pores That Cause ATP Leakage. mBio 9.

31. Lobato-Marquez D, Moreno-Cordoba I, Figueroa V, Diaz-Orejas R, Garcia-del Portillo F. 2015. Distinct type I and type II toxin-antitoxin modules control Salmonella lifestyle inside eukaryotic cells. Sci Rep 5:9374.

32. Han K, Kim KS, Bak G, Park H, Lee Y. 2010. Recognition and discrimination of target mRNAs by Sib RNAs, a cis-encoded sRNA family. Nucleic Acids Res 38:5851–66.

33. Mok WW, Patel NH, Li Y. 2010. Decoding toxicity: deducing the sequence requirements of IbsC, a type I toxin in Escherichia coli. J Biol Chem 285:41627–36.

34. Mirdita M, Schütze K, Moriwaki Y, Heo L, Ovchinnikov S, Steinegger M. 2022. ColabFold: making protein folding accessible to all. Nature Methods 19:679–682.

35. Dorr T, Lewis K, Vulic M. 2009. SOS response induces persistence to fluoroquinolones in Escherichia coli. PLoS Genet 5:e1000760.

36. Dorr T, Vulic M, Lewis K. 2010. Ciprofloxacin causes persister formation by inducing the TisB toxin in Escherichia coli. PLoS Biol 8:e1000317.

37. Wilmaerts D, Dewachter L, De Loose PJ, Bollen C, Verstraeten N, Michiels J. 2019. HokB Monomerization and Membrane Repolarization Control Persister Awakening. Mol Cell 75:1031–1042 e4.

38. Battesti A, Bouveret E. 2012. The bacterial two-hybrid system based on adenylate cyclase reconstitution in Escherichia coli. Methods 58:325–34.

39. Walker JE. 2013. The ATP synthase: the understood, the uncertain and the unknown. Biochem Soc Trans 41:1–16.

40. Vik SB, Antonio BJ. 1994. A mechanism of proton translocation by F1F0 ATP synthases suggested by double mutants of the a subunit. J Biol Chem 269:30364–9.

41. Kühlbrandt W. 2019. Structure and Mechanisms of F-Type ATP Synthases. Annu Rev Biochem 88:515–549.

42. Sobti M, Walshe JL, Wu D, Ishmukhametov R, Zeng YC, Robinson CV, Berry RM, Stewart AG. 2020. Cryo-EM structures provide insight into how E. coli F(1)F(o) ATP synthase accommodates symmetry mismatch. Nat Commun 11:2615.

43. Lee EJ, Pontes MH, Groisman EA. 2013. A bacterial virulence protein promotes pathogenicity by inhibiting the bacterium’s own F1Fo ATP synthase. Cell 154:146–56.

44. Lee EJ, Groisman EA. 2012. Control of a Salmonella virulence locus by an ATP-sensing leader messenger RNA. Nature 486:271–5.

45. Goormaghtigh F, Fraikin N, Putrins M, Hallaert T, Hauryliuk V, Garcia-Pino A, Sjodin A, Kasvandik S, Udekwu K, Tenson T, Kaldalu N, Van Melderen L. 2018. Reassessing the Role of Type II Toxin-Antitoxin Systems in Formation of Escherichia coli Type II Persister Cells. mBio 9.

46. Pontes MH, Lee EJ, Choi J, Groisman EA. 2015. Salmonella promotes virulence by repressing cellulose production. Proc Natl Acad Sci U S A 112:5183–8.

47. Fields PI, Swanson RV, Haidaris CG, Heffron F. 1986. Mutants of *Salmonella typhimurium* that cannot survive within the macrophage are avirulent. Proc Natl Acad Sci U S A 83:5189–93.

48. Datsenko KA, Wanner BL. 2000. One-step inactivation of chromosomal genes in *Escherichia coli* K-12 using PCR products. Proc Natl Acad Sci U S A 97:6640–5.

49. Davis RW, Bolstein D, Roth JR. 1980. Advanced Bacterial Genetics. Cold Spring Harbor Lab, Cold Spring Harbor, NY.

50. Snavely MD, Miller CG, Maguire ME. 1991. The *mgtB* Mg2+ transport locus of *Salmonella typhimurium* encodes a P-type ATPase. J Biol Chem 266:815–23.

51. Karimova G, Pidoux J, Ullmann A, Ladant D. 1998. A bacterial two-hybrid system based on a reconstituted signal transduction pathway. Proc Natl Acad Sci U S A 95:5752–6.

52. Maloy SR, Nunn WD. 1981. Selection for loss of tetracycline resistance by *Escherichia coli*. J Bacteriol 145:1110–1.

53. Rang C, Alix E, Felix C, Heitz A, Tasse L, Blanc-Potard AB. 2007. Dual role of the MgtC virulence factor in host and non-host environments. Mol Microbiol 63:605–22.

54. Choi S, Choi E, Cho YJ, Nam D, Lee J, Lee EJ. 2019. The Salmonella virulence protein MgtC promotes phosphate uptake inside macrophages. Nat Commun 10:3326.

55. Ansong C, Yoon H, Porwollik S, Mottaz-Brewer H, Petritis BO, Jaitly N, Adkins JN, McClelland M, Heffron F, Smith RD. 2009. Global systems-level analysis of Hfq and SmpB deletion mutants in Salmonella: implications for virulence and global protein translation. PLoS One 4:e4809.

56. Cho BK, Knight EM, Palsson BO. 2006. PCR-based tandem epitope tagging system for Escherichia coli genome engineering. Biotechniques 40:67–72.

57. Lee EJ, Groisman EA. 2010. An antisense RNA that governs the expression kinetics of a multifunctional virulence gene. Mol Microbiol 76:1020–33.

