## Supporting information for "A *Salmonella* type I toxin promotes systemic infection by inhibiting F_o_F_1_ ATP synthase"

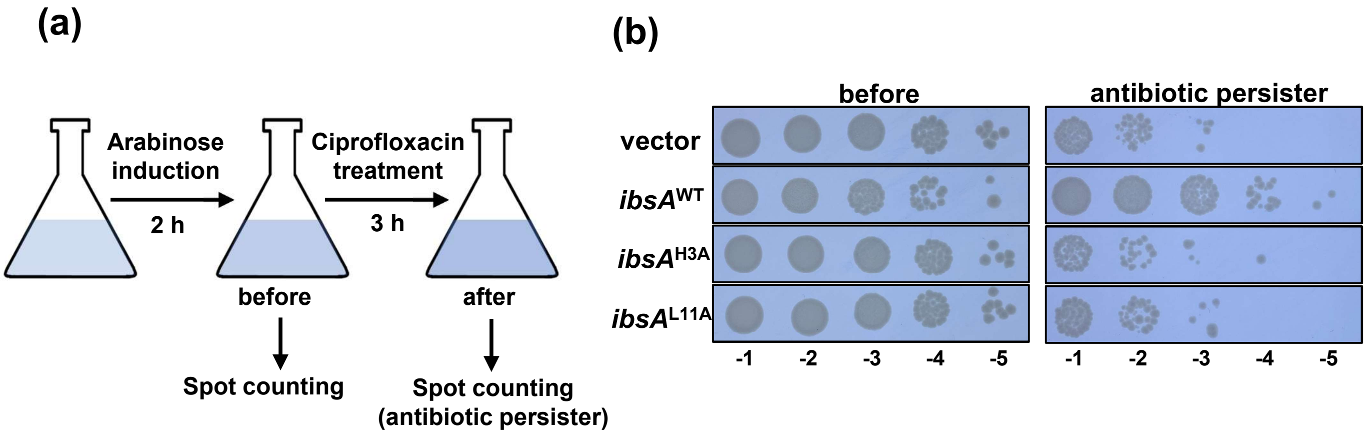


**Fig. S1. IbsA toxin promotes antibiotic-induced persister formation**

**(a)** Schematic cartoon of antibiotic-induced non-replicating cells analysis. **(b)** Ciprofloxacin-induced persister formation in *Salmonella* strains harboring an arabinose-inducible plasmid with the wild-type *ibsA*, *ibsA*$^{H3A}$, or *ibsA*$^{L11A}$ genes, and the empty vector. The strains were grown at 37°C in N-minimal medium containing 10 mM $Mg^{2+}$, and the expression of *ibsA* was induced by adding 10 mM arabinose when the cells reached an $OD_{600}$ of 0.1. After adding arabinose, the cells were grown for 2 hours and normalized by measuring $OD_{600}$. The cultures were then treated with 0.1 µg/ml ciprofloxacin and grown for an 3 hours. Cells harvested at indicated times were serial diluted and spotted on N-minimal agar plate containing 10 mM $Mg^{2+}$. The plates were incubated at 37°C for 18 hours.

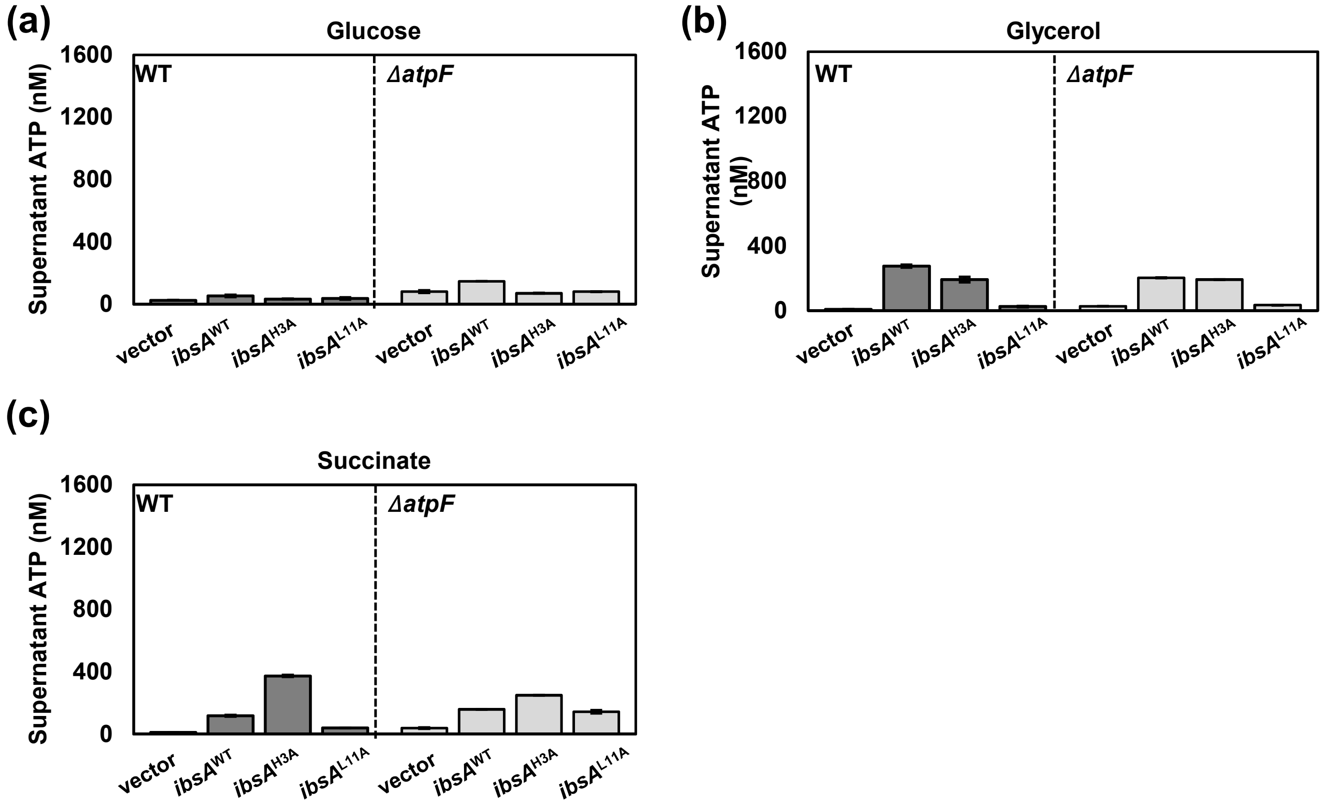


**Fig. S2. Extracellular ATP levels do not correlate with IbsA toxin activity**
**(a-c)** Supernatant ATP levels of 14028s or *ΔatpF Salmonella* in N-minimal medium containing 0.2% glucose **(a)**, 0.2% glycerol **(b)**, or 0.2% succinate **(c)** as described in Figure 3.

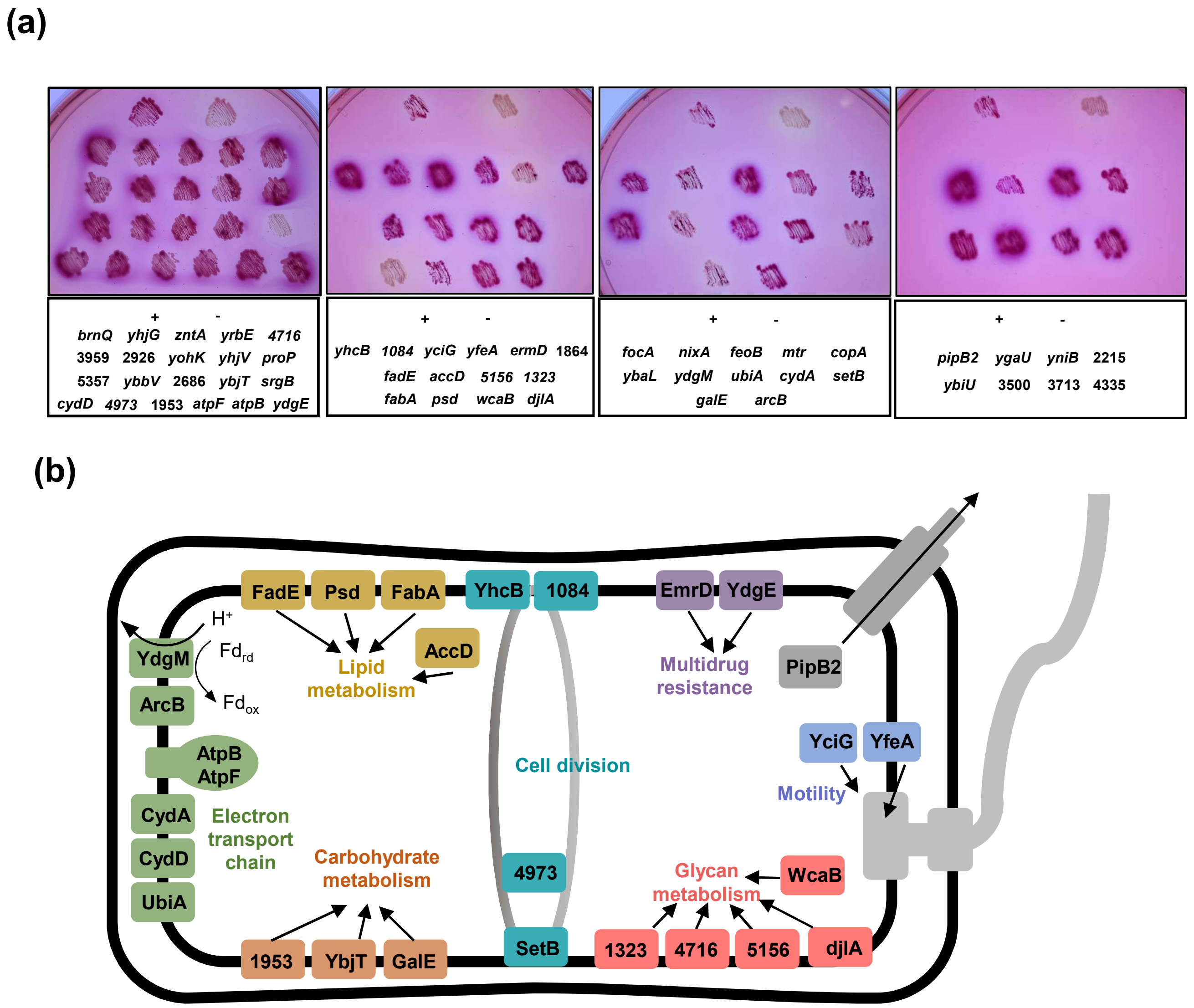


**Fig. S3. Bacterial two-hybrid screening identifies IbsA-interacting targets (a)** Images of selected colonies patched on MacConkey agar plates. The red-colored patches indicate a positive interaction with IbsA. In each clone, the genes fused in-frame to the T18 domain are indicated below the image. '+' indicates positive control, and '-' indicates a negative control (pUT18 and pKT25 empty vectors). **(b)** An illustration showing the location of isolated IbsA-interacting proteins in the cell. The colors represent the functional categories of these proteins, related to Table 1. Green, energy metabolism with oxidative phosphorylation; orange, carbohydrate metabolism; yellow, lipid metabolism; pink, glycan metabolism; teal, cell division; purple, multidrug resistance; blue, motility; gray, pathogenesis.

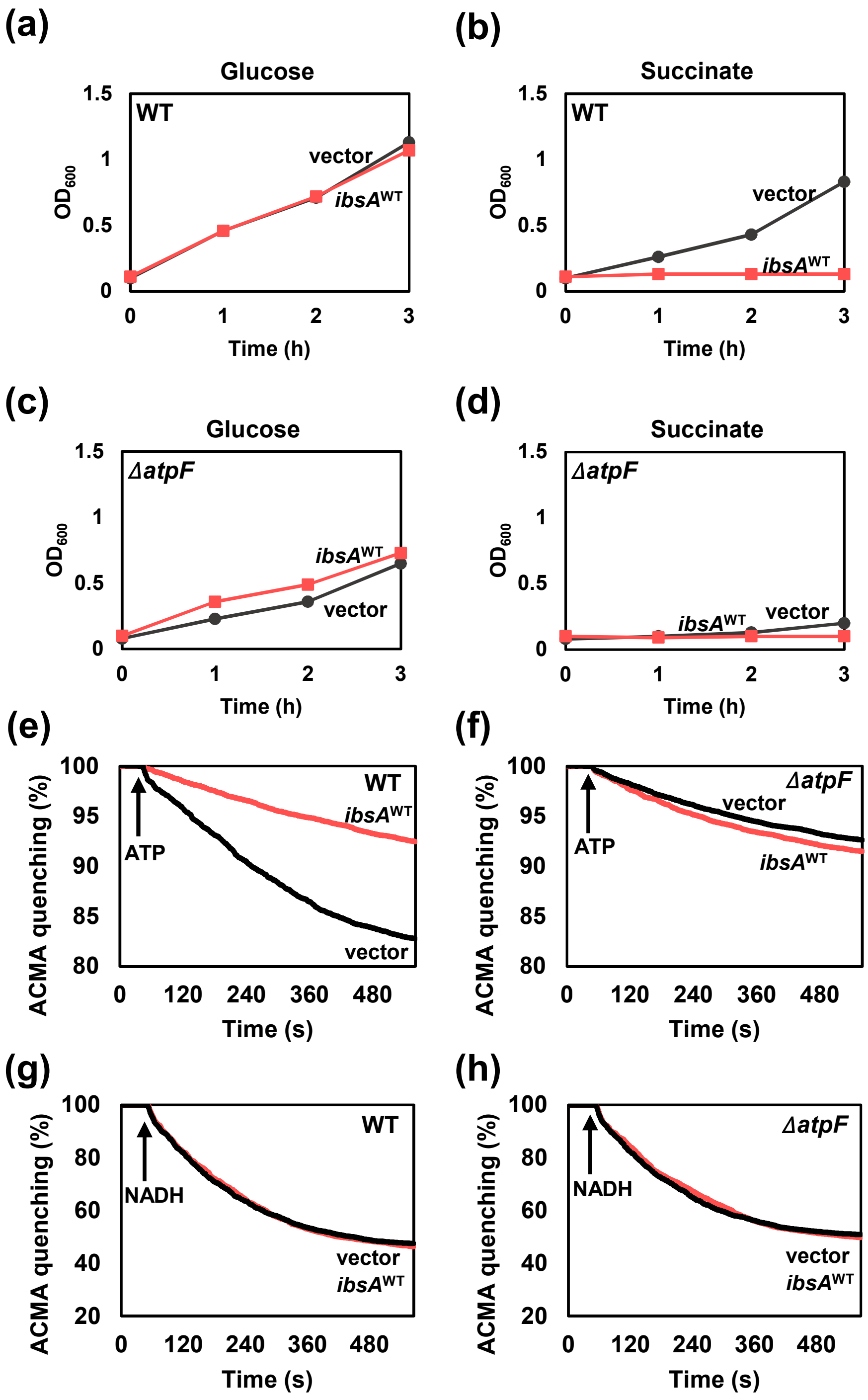


**Fig. S4. Growth inhibitory effect of IbsA is AtpF-dependent**

**(a-d)** Growth curves of wild-type **(a, b)** or *ΔatpF* **(c, d)** *Salmonella* strains harboring either the empty vector or pBAD33-*ibsA*. The strains were grown at 37°C in N-minimal medium containing 0.2% glucose **(a, c)** or 0.2% succinate **(b, d)** as the sole carbon sources. *ibsA* expression was induced by adding 10 mM arabinose when the cells reached at $OD_{600}$ of 0.1. The cells were grown for additional 2 hours and harvested for measurement of ATP levels in Figure 3. The absorbance at $OD_{600}$ was measured every hour. **(e-h)** Fluorescence quenching of ACMA in inverted membrane vesicles prepared from **(e)** wild-type *Salmonella* harboring either empty vector or pBAD33-*ibsA* and **(f)** *atpF* mutant *Salmonella* harboring either empty vector or pBAD33-*ibsA*. The reaction was initiated by adding 1 mM ATP. **(g-h)** Fluorescence quenching of ACMA in inverted membrane vesicles prepared as in **(e-f)**, with the reaction was initiated by adding 0.5 mM NADH.

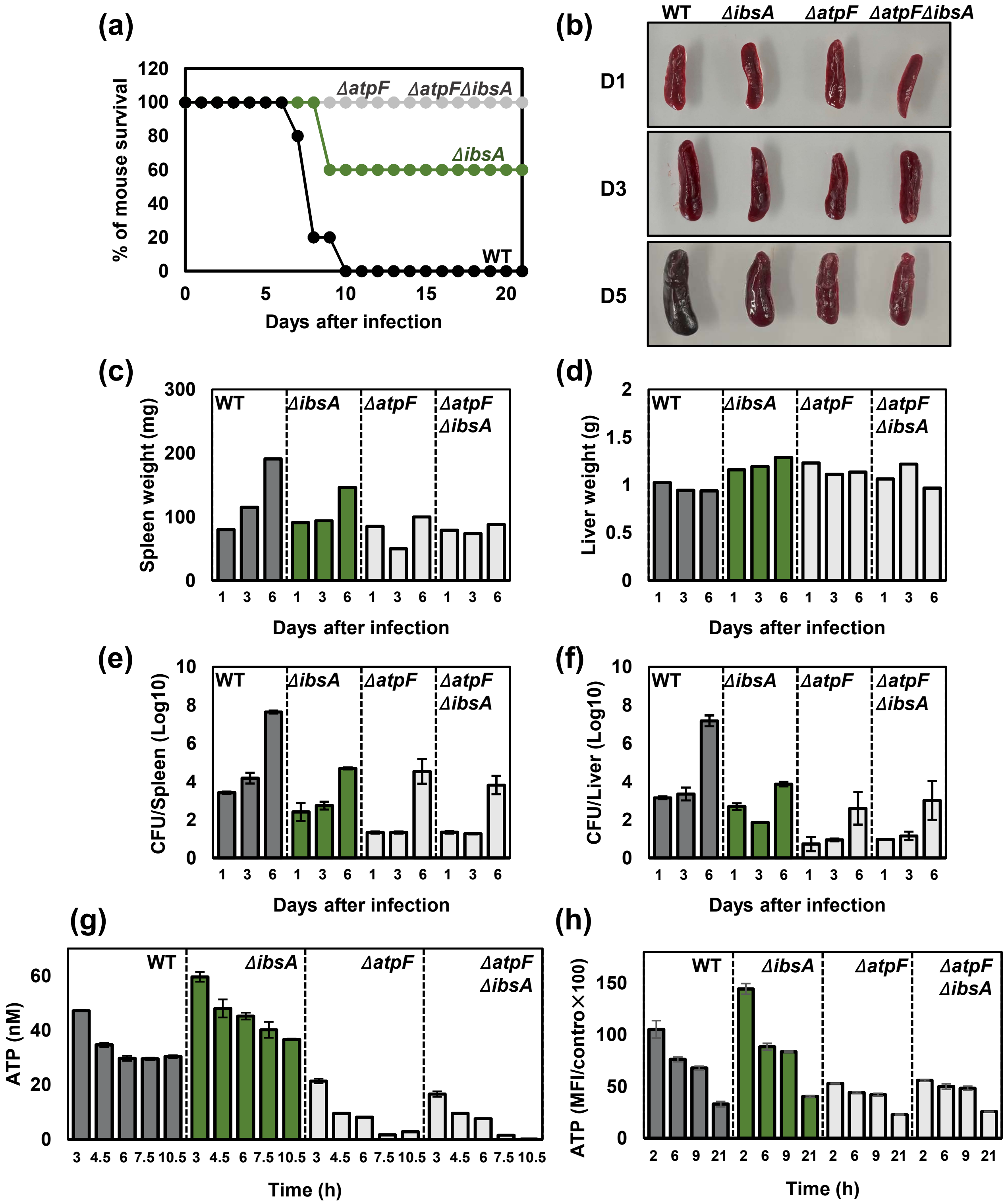
(a)
% of mouse survival
Days after infection
ΔatpF
ΔatpFΔibsA
ΔibsA
WT
(b)
WT
ΔibsA
ΔatpF
ΔatpFΔibsA
D1
D3
D5
(c)
Spleen weight (mg)
Days after infection
(d)
Liver weight (g)
Days after infection
(e)
CFU/Spleen (Log10)
Days after infection
(f)
CFU/Liver (Log10)
Days after infection
(g)
ATP (nM)
Time (h)
(h)
ATP (MFI/contro×100)
Time (h)

**Fig. S5. IbsA promotes mouse virulence and the effect is *atpF*-dependent**

**(a)** Survival of C3H/HeN mice inoculated intraperitoneally with approximately $10^3$ colony-forming units of wild-type, *ibsA* deletion, *atpF* deletion, and *atpF ibsA* double deletion *Salmonella* strains. The data are representative of two independent experiments, which gave similar results. **(b)** Appearance and relative size of spleens isolated from the mouse groups that received the above *Salmonella* strains at 1 (D1), 3 (D3), and 5 (D5) days post-infection. Shown are representative spleen samples from three independent experiments. **(c, d)** Spleen (c) and liver (d) weights of the mouse groups at 1, 3, and 5 days post-infection. Scale bar: 1 cm. **(e, f)** Bacterial loads in *Salmonella*-infected spleens (e) and livers (f). Mouse spleens and livers were harvested at days 1, 3, and 5. Organs were homogenized in PBS containing 0.1% Triton X-100. To measure the number of bacteria, homogenates were plated on LB solid media with appropriate dilutions. The numbers of *Salmonella* were determined and represented as $Log_{10}$ CFU/g of organ. **(g)** Temporal changes in intracellular ATP levels of the wild-type, *ibsA* deletion, *atpF* deletion, and *atpF ibsA* double deletion *Salmonella* strains, as described in Figure 9b. **(h)** Temporal changes in intracellular ATP levels of the wild-type, *ibsA* deletion, *atpF* deletion, and *atpF ibsA* double deletion *Salmonella* strains inside macrophages at 2, 6, 9, and 21 h post-infection, as described in Figure 9e.

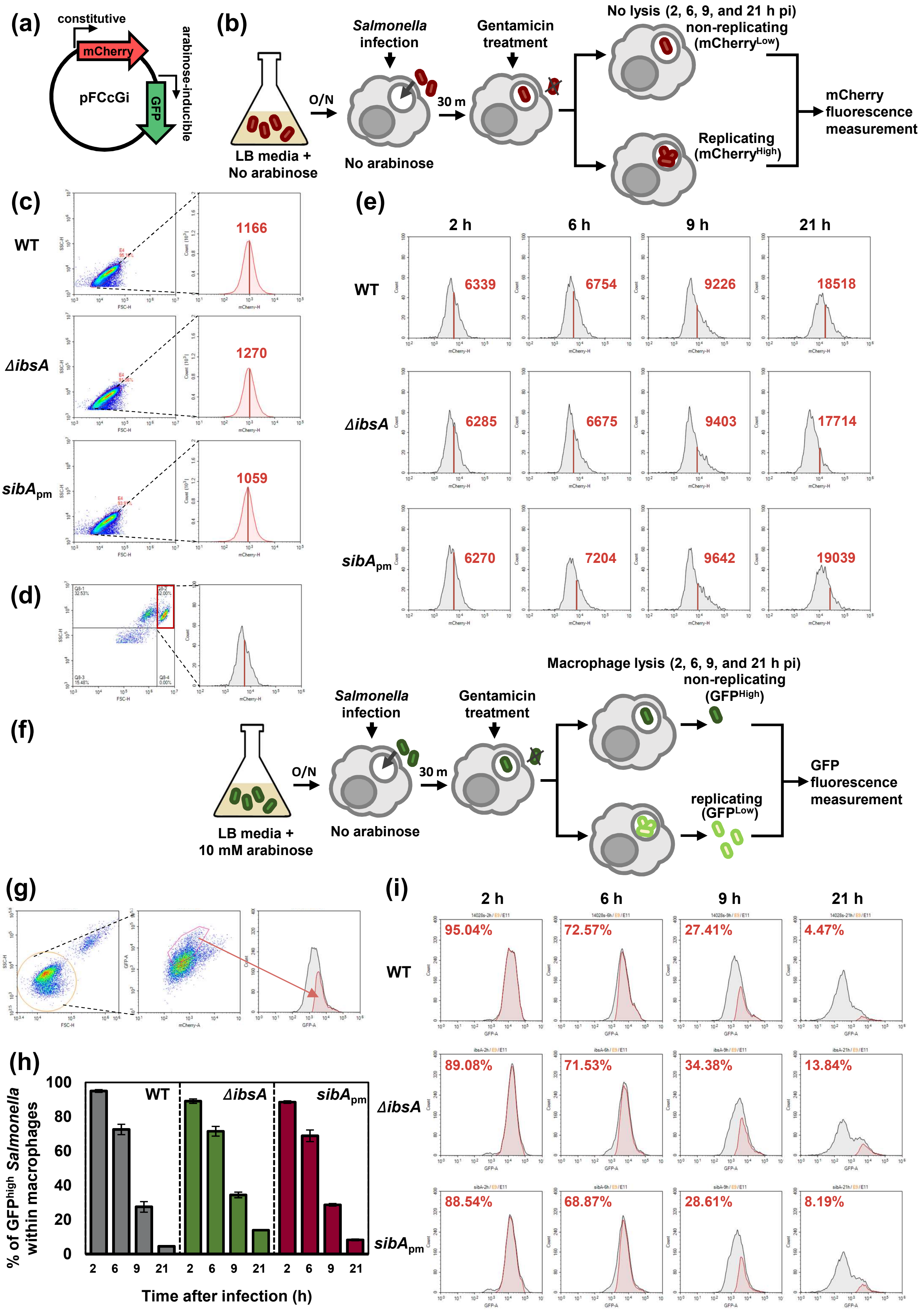

(a)
constitutive
mCherry
pFCcGi
GFP
arabinose-inducible
(b)
LB media +
No arabinose
O/N
Salmonella infection
No arabinose
30 m
Gentamicin treatment
No lysis (2, 6, 9, and 21 h pi)
non-replicating (mCherryLow)
Replicating (mCherryHigh)
mCherry fluorescence measurement
(c)
WT
1166
ΔibsA
1270
sibApm
1059
(d)
(e)
2 h
6 h
9 h
21 h
WT
6339
6754
9226
18518
ΔibsA
6285
6675
9403
17714
sibApm
6270
7204
9642
19039
(f)
LB media +
10 mM arabinose
O/N
Salmonella infection
No arabinose
30 m
Gentamicin treatment
Macrophage lysis (2, 6, 9, and 21 h pi)
non-replicating (GFPHigh)
replicating (GFPLow)
GFP fluorescence measurement
(g)
(h)
% of GFPhigh Salmonella within macrophages
Time after infection (h)
WT
ΔibsA
sibApm
(i)
2 h
6 h
9 h
21 h
WT
95.04%
72.57%
27.41%
4.47%
ΔibsA
89.08%
71.53%
34.38%
13.84%
sibApm
88.54%
68.87%
28.61%
8.19%

**Fig. S6. IbsA promotes intramacrophage survival in low multiplicity of infection**
**(a)** Schematic cartoon of pFCcGi dual fluorescence reporter. **(b)** Schematic representation of the intramacrophage survival assay using mCherry-expressing *Salmonella* strains listed in Fig. 8. Gentamicin protection assay of mCherry-expressing *Salmonella* was performed to determine replication efficiency within macrophages and mCherry fluorescence was monitored without macrophage lysis. **(c, d)** Gating strategy for flow cytometry analysis. **(c)** For the control, overnight-grown bacterial cells were gated using an FSC/SSC dot plot to select live bacteria. Then, mCherry plots were analyzed to determine the mean fluorescence intensities (MFIs) of the strains. Red vertical lines and numbers correspond to the MFIs of each strain. **(d)** For *Salmonella*-infected macrophages, cells were gated using an FSC/SSC dot plot to select macrophage cells (red rectangle), and mCherry plots were analyzed to determine the MFIs of *Salmonella* strains within macrophages. **(e)** Histograms of *Salmonella* strains expressing mCherry inside J774A.1 macrophages. Histograms indicate the mCherry fluorescence intensities of strains listed in Figure 8 at 2, 6, 9, and 21 h post-infection. Red vertical lines and numbers correspond to the MFIs of each strain. **(f)** Schematic representation of the GFP fluorescence dilution assay to analyze non-growing cells within macrophages. **(g)** Gating strategy for flow cytometry analysis. Cells were gated with FSC/SSC dot plot to select live bacteria. Then, mCherry/GFP plots were analyzed to select bacteria that emit high levels of green fluorescence, as shown in the grid. **(h)** The percentage of cells expressing high levels of GFP ($GFP^{High}$) was calculated from *Salmonella* strains harboring pFCcGi plasmid (n = 30,000 cells) at 2, 6, 9, and 21 h post-infection. Shown are the means and SD from three independent infections. **(i)** Histograms of *Salmonella* strains expressing GFP inside J774A.1 macrophages. Histograms indicate the GFP intensities of strains listed in Figure 8 at 2, 6, 9, and 21 h post-infection, with gates showing the fraction of the population exhibiting high and low GFP levels. Non-replicating $GFP^{High}$ cells are indicated as red histograms.

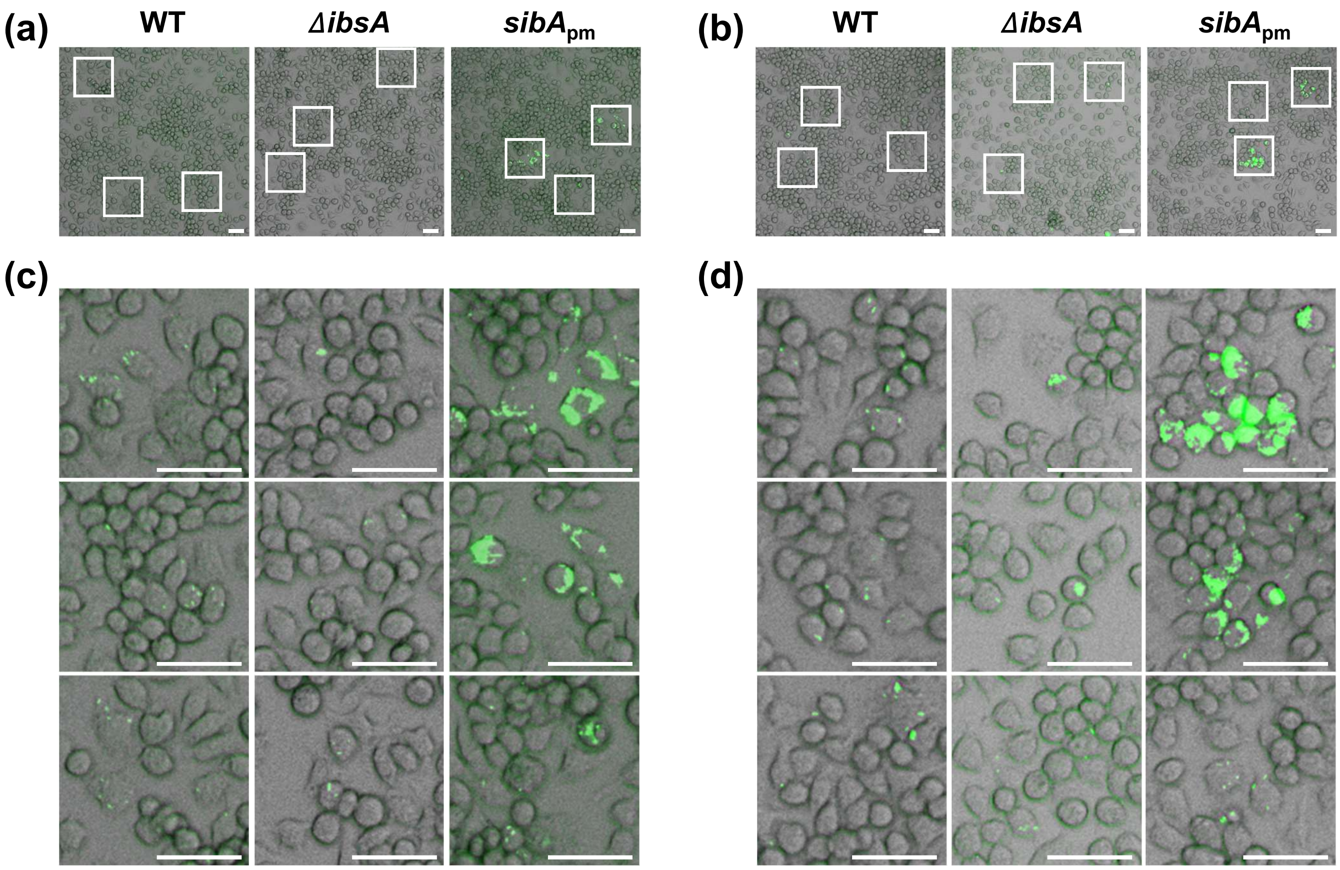


**Fig. S7. The *ibsA* mutant exhibits a defect in *Salmonella* spreading between macrophages**

**(a, b)** Two independent microscopic images of wild-type, *ΔibsA,* and *sibA*$_{pm}$ *Salmonella* strains expressing GFP inside J774A.1 cell lines at 21 h post-infection. J774A.1 cells were infected with *Salmonella* strains at a multiplicity of infection (MOI) of 0.1. The cells were incubated for 1 h to allow bacterial invasion and images were obtained at 21 h post-infection. Each grid corresponds to regions magnified in (c, d). **(c, d)** Magnified images from (a, b). Scale bar: 50 μm.

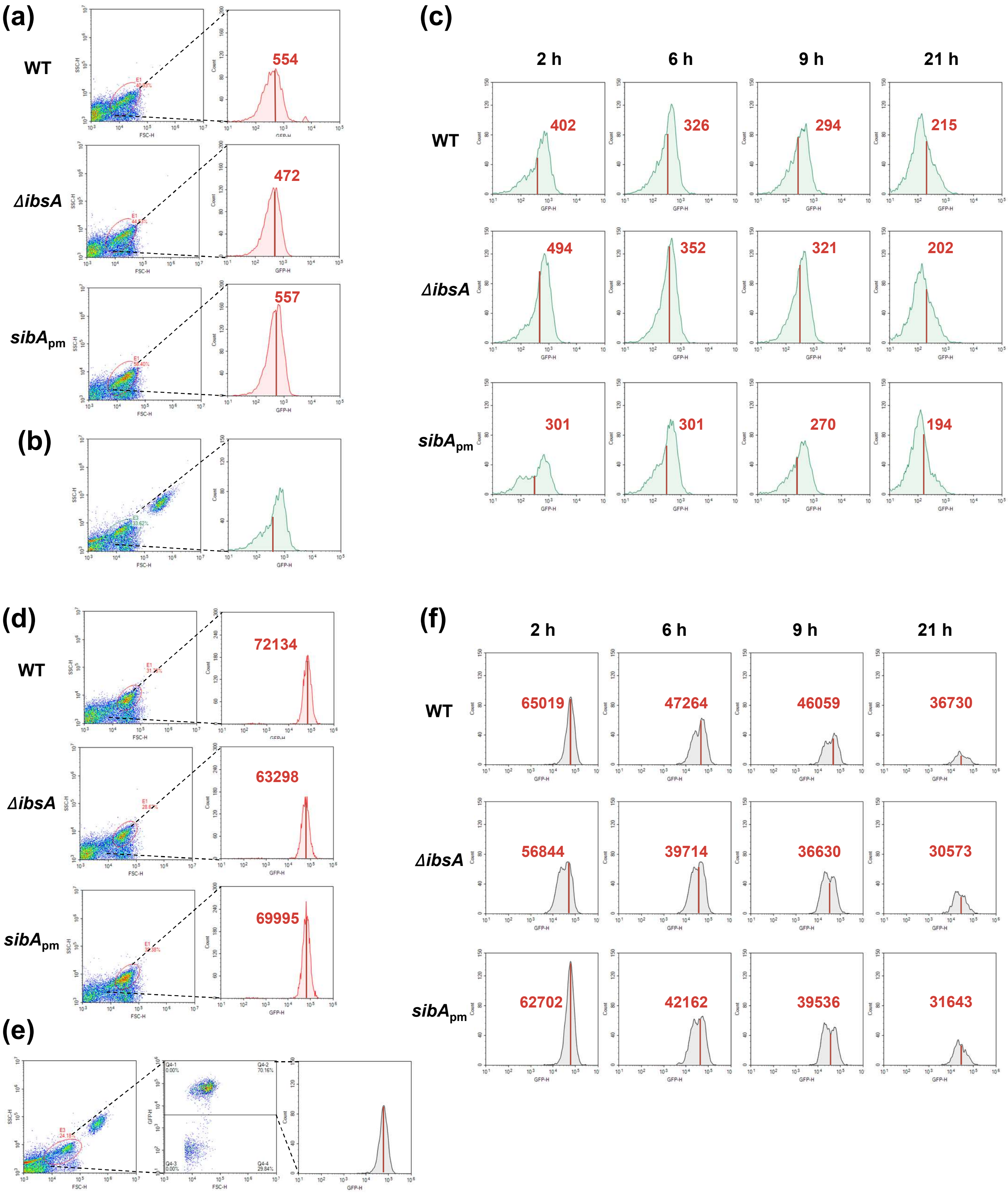
(a)
WT
554
ΔibsA
472
sibApm
557
(b)
(c)
2 h
6 h
9 h
21 h
WT
402
326
294
215
ΔibsA
494
352
321
202
sibApm
301
301
270
194
(d)
WT
72134
ΔibsA
63298
sibApm
69995
(e)
(f)
2 h
6 h
9 h
21 h
WT
65019
47264
46059
36730
ΔibsA
56844
39714
36630
30573
sibApm
62702
42162
39536
31643

**Fig. S8. Bacterial ATP levels inside macrophages can be measured using an ATP-responding GFP reporter**

**(a, b)** Gating strategy for flow cytometry analysis using $p_{lac}$303GFP. **(a)** For the control, overnight-grown bacterial cells were gated using an FSC/SSC dot plot to select live bacteria. Then, GFP plots were analyzed to determine the mean fluorescence intensities (MFIs) of the strains. Red vertical lines and numbers correspond to the MFIs of each strain. **(b)** For *Salmonella* strains inside macrophages, infected cells were lysed and gated using an FSC/SSC dot plot to select live bacterial cells (green circle), and GFP plots were analyzed to determine the MFIs of *Salmonella* strains within macrophages. **(c)** Histograms of *Salmonella* strains expressing GFP inside J774A.1 macrophages. Histograms indicate the fluorescence intensities of strains listed in Figure 9 at 2, 6, 9, and 21 h post-infection. Red vertical lines and numbers indicate the MFIs of each strain. **(d, e)** Gating strategy for flow cytometry analysis of the $p_{lac}$GFP control plasmid. **(d)** For the control, overnight-grown bacterial cells were gated using an FSC/SSC dot plot to select live bacteria. Then, GFP plots were analyzed to determine the mean fluorescence intensities (MFIs) of the strains. Red vertical lines and numbers correspond to the MFIs of each strain. **(e)** For *Salmonella* strains inside macrophages, infected cells were lysed and gated using an FSC/SSC dot plot to select bacterial cells (red circle), followed by a second gating (FSC-H versus GFP) to select bacteria emitting high levels of green fluorescence. Then, GFP plots were analyzed to determine the MFIs of *Salmonella* strains within macrophages. **(f)** Histograms of *Salmonella* strains expressing GFP inside J774A.1 macrophages. Histograms indicate the fluorescence intensities of strains listed in Figure 9 at 2, 6, 9, and 21 h post-infection. Red vertical lines and numbers correspond to the MFIs of each strain.

**Table S1. Bacterial strains and plasmids used in this study.**

| Name | Description | Reference |
|---|---|---|
| ***S. enterica* serovar Typhimurium** | | |
| 14028s | wild-type | (1) |
| MS7953s | *phoP7953*::Tn10 | (1) |
| SY100 | *ibsA/sibA::tetRA* | This study |
| SY51 | *ibsA* | This study |
| SY246 | *ibsA* $^{\text{His 3 Ala}}$ | This study |
| SY247 | *ibsA* $^{\text{Leu 11 Ala}}$ | This study |
| SY168 | *sibA*$_{\text{pm}}$ | This study |
| SY202 | *atpF* | This study |
| SY447 | *atpF, ibsA* | This study |
| SY329 | *ibsA*-8×Myc | This study |
| SY203 | *atpF*-2×HA | This study |
| SY42 | 14028s/pBAD33 | This study |
| SY60 | 14028s/pBAD33-*ibsA* | This study |
| SY70 | 14028s/pBAD33-*ibsA* $^{\text{Met 2 Ala}}$ | This study |
| SY71 | 14028s/pBAD33-*ibsA* $^{\text{His 3 Ala}}$ | This study |
| SY72 | 14028s/pBAD33-*ibsA* $^{\text{Gln 4 Ala}}$ | This study |
| SY84 | 14028s/pBAD33-*ibsA* $^{\text{Val 5 Ala}}$ | This study |
| SY85 | 14028s/pBAD33-*ibsA* $^{\text{Ile 6 Ala}}$ | This study |
| SY86 | 14028s/pBAD33-*ibsA* $^{\text{Ile 7 Ala}}$ | This study |
| SY87 | 14028s/pBAD33-*ibsA* $^{\text{Leu 8 Ala}}$ | This study |
| SY88 | 14028s/pBAD33-*ibsA* $^{\text{Ile 9 Ala}}$ | This study |
| SY89 | 14028s/pBAD33-*ibsA* $^{\text{Val 10 Ala}}$ | This study |
| SY90 | 14028s/pBAD33-*ibsA* $^{\text{Leu 11 Ala}}$ | This study |
| SY170 | 14028s/pBAD33-*ibsA* $^{\text{Leu 12 Ala}}$ | This study |
| SY91 | 14028s/pBAD33-*ibsA* $^{\text{Leu 13 Ala}}$ | This study |
| SY171 | 14028s/pBAD33-*ibsA* $^{\text{Ile 14 Ala}}$ | This study |
| SY73 | 14028s/pBAD33-*ibsA* $^{\text{Ser 15 Ala}}$ | This study |
| SY92 | 14028s/pBAD33-*ibsA* $^{\text{Phe 16 Ala}}$ | This study |
| SY74 | 14028s/pBAD33-*ibsA* $^{\text{Tyr 19 Ala}}$ | This study |
| SY350 | 14028s/pBAD33-*ibsA*$^{\text{GTG-started}}$-8×Myc | This study |
| SY351 | 14028s/pBAD33-*ibsA*$^{\text{ATG-started}}$-8×Myc | This study |
| SY435 | 14028s/pBAD33-*ibsA*$^{\text{GTG>TAG}}$ | This study |
| SY436 | 14028s/pBAD33-*ibsA*$^{\text{ATG>GCG}}$ | This study |
| SY209 | *atpF*/pBAD33 | This study |
| SY210 | *atpF*/pBAD33-*ibsA* | This study |
| SY209 | *atpF*/pBAD33-*ibsA* $^{\text{His 3 Ala}}$ | This study |
| SY210 | *atpF*/pBAD33-*ibsA* $^{\text{Leu 11 Ala}}$ | This study |
| SY117 | 14028s/ pBAD33-GFP-*ibsA* | This study |
| SY312 | 14028s/ pBAD33-GFP-*ibsA* $^{\text{His 3 Ala}}$ | This study |
| SY313 | 14028s/ pBAD33-GFP-*ibsA* $^{\text{Leu 11 Ala}}$ | This study |
| SY213 | *atpF*-HA, pBAD33 | This study |
| SY214 | *atpF*-HA, pBAD33-GFP-*ibsA* | This study |
| SY437 | *atpF*-HA, pBAD33-GFP-*ibsA* $^{\text{His 3 Ala}}$ | This study |
| SY438 | *atpF*-HA, pBAD33-GFP-*ibsA* $^{\text{Leu 11 Ala}}$ | This study |
| SY169 | 14028s/pFCcGi | This study |
| SY294 | *ibsA*/pFCcGi | This study |
| SY297 | *sibA*$_{\text{pm}}$/pFCcGi | This study |
| SY344 | 14028s/pFPV25.1 | This study |
| SY420 | *ibsA*/pFPV25.1 | This study |

| | | |
|---|---|---|
| SY421 | *sibA*$_{pm}$/pFPV25.1 | This study |
| SY455 | 14028s/plac303GFP | This study |
| SY456 | *ibsA*/plac303GFP | This study |
| SY457 | *sibA*$_{pm}$/plac303GFP | This study |
| SY458 | *atpF*/plac303GFP | This study |
| SY459 | *atpFibsA*/plac303GFP | This study |
| ***Escherichia coli*** | | |
| DH5α | *fhuA2 lac(del)U169 phoA glnV44 Φ80' lacZ(del)M15 gyrA96 recA1 relA1 endA1 thi-1 hsdR17.* | (2) |
| SY57 | DH5α/pBAD33-*ibsA* | This study |
| SY65 | DH5α/pBAD33-*ibsA* $^{\text{Met 2 Ala}}$ | This study |
| SY66 | DH5α/pBAD33-*ibsA* $^{\text{His 3 Ala}}$ | This study |
| SY67 | DH5α/pBAD33-*ibsA* $^{\text{Gln 4 Ala}}$ | This study |
| SY75 | DH5α/pBAD33-*ibsA* $^{\text{Val 5 Ala}}$ | This study |
| SY76 | DH5α/pBAD33-*ibsA* $^{\text{Ile 6 Ala}}$ | This study |
| SY77 | DH5α/pBAD33-*ibsA* $^{\text{Ile 7 Ala}}$ | This study |
| SY78 | DH5α/pBAD33-*ibsA* $^{\text{Leu 8 Ala}}$ | This study |
| SY79 | DH5α/pBAD33-*ibsA* $^{\text{Ile 9 Ala}}$ | This study |
| SY80 | DH5α/pBAD33-*ibsA* $^{\text{Val 10 Ala}}$ | This study |
| SY81 | DH5α/pBAD33-*ibsA* $^{\text{Leu 11 Ala}}$ | This study |
| SY160 | DH5α/pBAD33-*ibsA* $^{\text{Leu 12 Ala}}$ | This study |
| SY82 | DH5α/pBAD33-*ibsA* $^{\text{Leu 13 Ala}}$ | This study |
| SY161 | DH5α/pBAD33-*ibsA* $^{\text{Ile 14 Ala}}$ | This study |
| SY68 | DH5α/pBAD33-*ibsA* $^{\text{Ser 15 Ala}}$ | This study |
| SY83 | DH5α/pBAD33-*ibsA* $^{\text{Phe 16 Ala}}$ | This study |
| SY69 | DH5α/pBAD33-*ibsA* $^{\text{Tyr 19 Ala}}$ | This study |
| SY348 | DH5α /pBAD33- *ibsA*$^{\text{GTG-started}}$-8×Myc | This study |
| SY349 | DH5α /pBAD33- *ibsA*$^{\text{ATG-started}}$-8×Myc | This study |
| SY433 | DH5α /pBAD33- *ibsA*$^{\text{GTG>TAG}}$ | This study |
| SY434 | DH5α /pBAD33- *ibsA*$^{\text{ATG>GCG}}$ | This study |
| ENC1343 | DH5α/pBAD33-GFP | This study |
| SY115 | DH5α/pBAD33-GFP-*ibsA* | This study |
| SY226 | DH5α/pBAD33-GFP-*ibsA* $^{\text{His 3 Ala}}$ | This study |
| SY227 | DH5α/pBAD33-GFP-*ibsA* $^{\text{Leu 11 Ala}}$ | This study |
| SY95 | DH5α/pKT25-*ibsA* | This study |
| SY124 | DH5α/pKT25-*ibsA* $^{\text{His 3 Ala}}$ | This study |
| SY162 | DH5α/pKT25-*ibsA* $^{\text{Leu 11 Ala}}$ | This study |
| SY218 | DH5α/pUT18-atpF | This study |
| BTH101 | F-, *cya-854*, *recA1*, *endA1*, *gyrA96* (Nal$^{r}$), *thi1*, *hsdR17*, *spoT1*, *rfbD1*, *glnV44(AS)* | (3) |
| LJ18 | BTH101/pUT18-*mgtC*, pKT25-*mgtR* | (4) |
| LJ27 | BTH101/pUT18-*mgtC*, pKT25 | (4) |
| SY219 | BTH101/pUT18-*atpF*, pKT25-*ibsA* | This study |
| SY220 | BTH101/pUT18-*atpF*, pKT25-*ibsA* $^{\text{His 3 Ala}}$ | This study |
| SY221 | BTH101/pUT18-*atpF*, pKT25-*ibsA* $^{\text{Leu 11 Ala}}$ | This study |
| **Plasmids** | | |
| pBAD33 | pACYC184 *ori* Cm$^{R}$ | (5) |
| pBAD33-GFP | pBAD33 *gfp* lacking the stop codon | This study |
| pKD3 | repR$_{6K\gamma}$ Ap$^{R}$ FRT Cm$^{R}$ FRT | (6) |
| pKD4 | repR$_{6K\gamma}$ Ap$^{R}$ FRT Km$^{R}$ FRT | (6) |
| pKD46 | rep$_{pSC101}$$^{ts}$ Ap$^{R}$ P$_{araBAD}$ γ β exo | (6) |
| pKD13-2×HA | pKD13 derivative harboring 2HA | (7) |

| | | |
|---|---|---|
| pBOP508 | $repR_{R6K}$ $Ap^R$ 8×myc FRT $Km^R$ FRT | (8) |
| pCP20 | $rep_{pSC101}{}^{ts}$ $Ap^R$ $Cm^R$ *cI857* $\lambda P_R$*flp* | (6) |
| pUT18 | $p_{lac}$ ColEI *ori* $Ap^R$ | (9) |
| pUT18c | $p_{lac}$ ColEI *ori* $Ap^R$ | (9) |
| pKT25 | $p_{lac}$ p15A*ori* $Km^R$ | (9) |
| pFPV25.1 | pfpv25 prpsM-gfpmut3 | (10) |
| pFCcGi | rpsM::mCherry and PBAD::gfpmut3a promoter fusions in pFPV25.1, $Ap^R$ | (11) |
| plac303GFP | pfpv25 $p_{lac1\text{-}6}$-*mgtC* leader 303-*gfp* | This study |

### Supplemental references


1. Fields PI, Swanson RV, Haidaris CG, Heffron F. 1986. Mutants of *Salmonella typhimurium* that cannot survive within the macrophage are avirulent. Proc Natl Acad Sci U S A 83:5189-93.
2. Taylor RG, Walker DC, McInnes RR. 1993. *E. coli* host strains significantly affect the quality of small scale plasmid DNA preparations used for sequencing. Nucleic Acids Res 21:1677-8.
3. Karimova G, Pidoux J, Ullmann A, Ladant D. 1998. A bacterial two-hybrid system based on a reconstituted signal transduction pathway. Proc Natl Acad Sci U S A 95:5752-6.
4. Choi S, Choi E, Cho YJ, Nam D, Lee J, Lee EJ. 2019. The Salmonella virulence protein MgtC promotes phosphate uptake inside macrophages. Nat Commun 10:3326.
5. Guzman LM, Belin D, Carson MJ, Beckwith J. 1995. Tight regulation, modulation, and high-level expression by vectors containing the arabinose PBAD promoter. J Bacteriol 177:4121-30.
6. Datsenko KA, Wanner BL. 2000. One-step inactivation of chromosomal genes in *Escherichia coli* K-12 using PCR products. Proc Natl Acad Sci U S A 97:6640-5.
7. Ansong C, Yoon H, Porwollik S, Mottaz-Brewer H, Petritis BO, Jaitly N, Adkins JN, McClelland M, Heffron F, Smith RD. 2009. Global systems-level analysis of Hfq and SmpB deletion mutants in Salmonella: implications for virulence and global protein translation. PLoS One 4:e4809.
8. Cho BK, Knight EM, Palsson BO. 2006. PCR-based tandem epitope tagging system for Escherichia coli genome engineering. Biotechniques 40:67-72.
9. Karimova G, Ullmann A, Ladant D. 2001. Protein-protein interaction between *Bacillus stearothermophilus* tyrosyl-tRNA synthetase subdomains revealed by a bacterial two-hybrid system. J Mol Microbiol Biotechnol 3:73-82.
10. Valdivia RH, Falkow S. 1996. Bacterial genetics by flow cytometry: rapid isolation of Salmonella typhimurium acid-inducible promoters by differential fluorescence induction. Mol Microbiol 22:367-78.
11. Figueira R, Watson KG, Holden DW, Helaine S. 2013. Identification of salmonella pathogenicity island-2 type III secretion system effectors involved in intramacrophage replication of S. enterica serovar typhimurium: implications for rational vaccine design. mBio 4:e00065.

**Table S2. Primers used in this study.**

| Name | Sequence (from 5' to 3')* | |
|---|---|---|
| **Deletion or tagging** | | |
| SZ008 | CTTGTCTGTACAAAGCGTTATGACCAGCAGGCCATTTTGCTGTAGGCTGGAG<br>CTGCTTCG | *ibsA* deletion Km$^{R}$ cassette insertion |
| SZ009 | TAAGGGTGAGAGGGGATCTCTCCCCCCCTCTGATTGTCTGCATATGAATATC<br>CTCCTTAG | *ibsA* deletion Km$^{R}$ cassette insertion |
| SZ046 | CCCCCGCCATATCTCCTGTCCCTGGTCATAAGACCGGGGATTAAGACCCACT<br>TTCACATTTAAG | *ibsA* Tet$^{R}$ cassette insertion |
| SZ047 | TCAAAAAGATCCAGACCGAGAGCGAGAAGTAAAACCGCGTCTAAGCACTT<br>GTCTCCTGTTTAC | *ibsA* Tet$^{R}$ cassette insertion |
| SZ058 | TTCCTTTACCGGCGGTCGAAAAAAAAGCAGCCGGCCCCGCGAGAT | *sibA* promotor mutation |
| SZ059 | ATCTCGCGGGGCCGGCTGCTTTTTTTTCGACCGCCGGTAAAGGAA | *sibA* promotor mutation |
| SZ107 | GTTATGATGGCCCAGGTCATC | *ibsA* $^{H3A}$ substitution |
| SZ108 | GATGACCTGGGCCATCATAAC | *ibsA* $^{H3A}$ substitution |
| SZ109 | CTGATTGTTGCGTTACTGATA | *ibsA* $^{L11A}$ substitution |
| SZ110 | TATCAGTAACGCAACAATCAG | *ibsA* $^{L11A}$ substitution |
| SZ111 | GATCGATTCGCTGTTAAACG | *ibsA* amino acid mutation PCR forward |
| SZ112 | CTGGTAAAAGGCGGCGACTA | *ibsA* amino acid mutation PCR forward |
| SZ173 | CATACTGATTGTTCTGTTACTGATAAGTTTCGCAGCTTACATCGGATCCAGA<br>ATTCGTGAT | *ibsA*-8×Myc Km$^{R}$ cassette insertion |
| SZ174 | TAAGGGTGAGAGGGGATCTCTCCCCCCCTCTGATTGTCTGGAGCTCGATCCG<br>TCGACC | *ibsA*-8×Myc Km$^{R}$ cassette insertion |
| SZ098 | TGCTAACAGCGACATCGTGGATAAACTTGTCGCTGAACTGTATCCGTATGAT<br>GTTCCGGATTATGCA TAA TGTAGGCTGGAGCTGCTTCG | *atpF*-2×HA Km$^{R}$ cassette insertion |
| SZ099 | CGAGCTACCGTAACAAATTCAGACATCAGCCCCTCCCTCCCATATGAATATC<br>CTCCTTAG | *atpF* deletion Km$^{R}$ cassette insertion |
| SZ100 | GAGCAATATCAGAAGGTTAACTAGATAGAGGCATTGTGCTTGTAGGCTGGA<br>GCTGCTTCG | *atpF* deletion Km$^{R}$ cassette insertion |
| **Cloning** | | |
| SZ001 | GCGGTACCCCTGACCAGCAGGCCATTTTGCGTGGGCGATCCAGTAACGCA | *ibsA*$^{GTG\text{-}started}$-KpnI |
| SZ002 | CCCAAGCTTTTAGTAAGCTGCGAAACTTA | *ibsA*-HindIII |
| SZ003 | GCGGTACCCCGCTTTAATAAGGAAAGGGTTATGATGCACCAGGTCATCAT | *ibsA*-KpnI |
| SZ013 | GAAAGGGTTGCGGCGCACCAG | *ibsA*$^{ATG>GCG}$ |
| SZ014 | CTGGTGCGCCGCAACCCTTTC | *ibsA*$^{ATG>GCG}$ |
| SZ019 | GCGGTACCCCGCTTTAATAAGGAAAGGGTTATGGCGCACCAGGTC | *ibsA* $^{M2A}$ |
| SZ020 | GCGGTACCCCGCTTTAATAAGGAAAGGGTTATGATGGCCCAGGTCATC | *ibsA* $^{H3A}$-KpnI |
| SZ021 | GCGGTACCCCGCTTTAATAAGGAAAGGGTTATGATGCACGCGGTCATCATA | *ibsA* $^{Q4A}$-KpnI |
| SZ022 | GCGGTACCCCGCTTTAATAAGGAAAGGGTTATGATGCACCAGGCCATCATA<br>CTG | *ibsA* $^{V5A}$-KpnI |
| SZ023 | GCGGTACCCCGCTTTAATAAGGAAAGGGTTATGATGCACCAGGTCGCCATA<br>CTGATT | *ibsA* $^{I6A}$-KpnI |
| SZ024 | GCGGTACCCCGCTTTAATAAGGAAAGGGTTATGATGCACCAGGTCATCGCA<br>CTGATTGTT | *ibsA* $^{I7A}$-KpnI |
| SZ025 | GCGGTACCCCGCTTTAATAAGGAAAGGGTTATGATGCACCAGGTCATCATA<br>GCGATTGTTCTG | *ibsA* $^{L8A}$-KpnI |
| SZ026 | GCGGTACCCCGCTTTAATAAGGAAAGGGTTATGATGCACCAGGTCATCATA<br>CTGGCTGTTCTGTTA | *ibsA* $^{I9A}$-KpnI |
| SZ027 | GCGGTACCCCGCTTTAATAAGGAAAGGGTTATGATGCACCAGGTCATCATA<br>CTGATTGCTCTGTTACTG | *ibsA* $^{V10A}$-KpnI |
| SZ028 | CCCAAGCTTTTAGTAAGCTGCGAAACTTATCAGTAACGCAACAATCAG | *ibsA* $^{L11A}$-HindIII |
| SZ029 | CCCAAGCTTTTAGTAAGCTGCGAAACTTATCAGTGCCAGAACAAT | *ibsA* $^{L12A}$-HindIII |
| SZ030 | CCCAAGCTTTTAGTAAGCTGCGAAACTTATCGCTAACAGAAC | *ibsA* $^{L13A}$-HindIII |
| SZ031 | CCCAAGCTTTTAGTAAGCTGCGAAACTTGCCAGTAACAG | *ibsA* $^{I14A}$-HindIII |
| SZ032 | CCCAAGCTTTTAGTAAGCTGCGAAAGCTATCAGTAA | *ibsA* $^{S15A}$-HindIII |

| | | |
|---|---|---|
| SZ033 | CCCAAGCTTTTAGTAAGCTGCGGCACTTATCAG | *ibsA* $^{F16AS}$-HindIII |
| SZ034 | CCCAAGCTTTGTCTGTTAGGCAGCTGCGAAACTTA | *ibsA* $^{Y19A}$-HindIII |
| SZ007 | GCTCTAGAGATGCACCAGGTCATCATA | *ibsA*-XbaI |
| SZ006 | GGGGTACCCCGTAAGCTGCGAAACTTATCA | *ibsA*-KpnI |
| SZ070 | GCTCTAGAGATGATGGCCCAGGTCATC | *ibsA* $^{H3A}$-XbaI |
| SZ037 | TCTAGAGAATCTTAACGCAACAATCCT | *atpF*-XbaI |
| SZ038 | GGTACCCCCAGTTCAGCGACAAGTTTAT | *atpF*-KpnI |
| SZ068 | GCTCTAGAGCACCAGGTCATCATACTGAT | *ibsA*-XbaI (GFP) |
| SZ105 | GCTCTAGAGGCCCAGGTCATCATACTGAT | *ibsA* $^{H3A}$-XbaI (GFP) |
| SZ176 | GCGGTACCCCTGACCAGCAGGCCATTTTGCATG | *ibsA* $^{GTG\text{-}started}$-8×Myc-KpnI |
| SZ177 | GCGGTACCCCGCTTTAATAAGGAAAGGGTTATGATG | *ibsA* $^{ATG\text{-}started}$-8×Myc-KpnI |
| SZ178 | CCCAAGCTTTGTTTCGACGCGTATCTCGC | *ibsA*-8×Myc-HindIII |
| 9870 | cggaattcctttacactttaagctttttatgtttatgttgtgtggaCAAATTCATGCAGGAGTAATATG | pLac1-6_*mgtC*UTR1-23F_EcoRI |
| 8117 | AAGTCTAGATTAACATACGTTCCTCCATT | MgtC +302 R XbaI |
| KU234 | GGGGTACCCTTTAAGAAGGAGATATACAT | gfp-KpnI |
| KU238 | GCTCTAGAGCTTTGTATAGTTCATCCAT | gfp-Xba1 |
| **Northern blot** | | |
| SZ_myc | CAGATCTTCTTCGCTAATCAGTTTCTGTTC | myc northern |
| SZ_sibA | GGGGGAGAGATCCCCTCTCA | sibA northern |
| SZ_rrfD | CTACGGCGTTTCACTTCTGAGTTC | rrfD northern |
| **Quantitative real-time PCR** | | |
| SZ_ibsA-F | GGGCTGAAACGGGAAAGC | ibsA qRT-PCR |
| SZ_ibsA-R | GCATCATAACCCTTTCCTTATTAAAGC | ibsA qRT-PCR |
| SZ_rrsH-F | CCAGCAGCCGCGGTAAT | rrsH qRT-PCR |
| SZ_rrsH-R | TTTACGCCCAGTAATTCCGATT | rrsH qRT-PCR |
